# Rare coding variants in *NOX4* link high superoxide levels to psoriatic arthritis mutilans

**DOI:** 10.1101/2023.06.06.543425

**Authors:** Sailan Wang, Pernilla Nikamo, Leena Laasonen, Bjorn Gudbjornsson, Leif Ejstrup, Lars Iversen, Ulla Lindqvist, Jessica J. Alm, Jesper Eisfeldt, Xiaowei Zheng, Sergiu-Bogdan Catrina, Fulya Taylan, Raquel Vaz, Mona Ståhle, Isabel Tapia-Paez

## Abstract

Psoriatic arthritis mutilans (PAM) is the rarest and most severe form of psoriatic arthritis. PAM is characterized by erosions of the small joints of hands and feet and osteolysis leading to joint disruption. Despite its severity, the underlying mechanisms are unknown, and no candidate susceptibility genes have hitherto been identified. We aimed to investigate the genetic basis of PAM. We performed massive parallel sequencing of sixty-one patients’ genomes from the PAM Nordic cohort. We validated the rare variants found by Sanger sequencing and genotyped additional psoriasis, psoriatic arthritis, and control cohorts. We then tested the role of the variants using *in vivo* and *in vitro* models. We found rare variants with a minor allele frequency (MAF) below 0.0001 in the NADPH oxidase 4 (*NOX4*) in four patients. *In silico* predictions show that the identified variants are potentially damaging. NOXs are the only enzymes producing reactive oxygen species (ROS). ROS are highly reactive molecules important role in the regulation of signal transduction. NOX4 is specifically involved in the differentiation of osteoclasts, the cells implicated in bone resorption. Functional follow-up studies using cell culture, zebrafish models, and measurement of ROS in patients uncovered that the *NOX4* variants found in this study increase the levels of ROS both in *vitro* and *in vivo.* We propose *NOX4* as the first candidate susceptibility gene for PAM. Our study links high levels of ROS caused by *NOX4* variants to the development of PAM, opening the possibility for a potential therapeutic target.

## Introduction

Psoriasis is a common inflammatory skin disease characterized by an abnormal hyperproliferation of keratinocytes, activated dendritic cells and infiltration of T lymphocytes in lesions ^1^. The incidence of psoriasis in Europeans is ∼3%, and it has been estimated that ∼30% of psoriasis patients develop psoriasis arthritis (PsA), a systemic chronic inflammatory disease with clinical features such as arthritis, enthesitis, dactylitis, tendonitis and cutaneous psoriasis ^2^. PsA is often classified into five subtypes: distal interphalangeal predominant, asymmetric oligoarticular, symmetric polyarthritis, spondylitis, and psoriatic arthritis mutilans(PAM) ^3^. PAM is the rarest and most severe form of PsA and is characterized by the shortening of one or more digits, due to severe osteolysis of the bones, a deformity known as “digital telescoping” or “opera glass finger”. PAM patients suffer from severe joint destruction causing flail joints, and the progress of the deformities is rapid once the disease starts ^4, 5^. The skin phenotype in PAM patients is often described as mild ^6^. Even though the overall prevalence of PAM is uncertain, several case studies have reported on PAM patients in different populations ^5, 7–9^, and in a Nordic PAM study it was estimated to have a prevalence of 3.7 cases per million habitants ^6^. Clinical and radiographic details of patients in the Nordic PAM study have been described ^4, 6, 7, 10, 11^.

Genetic factors play an important role in the development of psoriasis and PsA, with dozens of susceptibility genes identified, and many but not all genetic signals overlapping ^2^. Most of the known susceptibility genes act via the HLA locus, IFN, NF-κB and IL23/17 signaling pathways, with some genes involved in skin barrier integrity such as *LCE3B-LCE3C* ^2^. It is also thought that genetic and environmental factors such as smoking, injuries and infections play a role in the etiology of psoriatic disorders ^12^. Humans have suffered from PAM since ancient times, skeletal remains with characteristic lesions of PAM have been found in a Byzantine monastery in Israel^13^. Today, PAM has been reported in many studies from all over the world, reviewed in ^14^, but the true prevalence of PAM is difficult to determine due to difficulties in clinical diagnosis and lack of biomarkers.

The nicotinamide adenine dinucleotide phosphate oxidase 4 (*NOX4*) gene (OMIM: 605261) encodes a protein that contains six transmembrane domains and in its cytosolic part, a flavin adenine dinucleotide (FAD) and a NADPH binding domain. NOX4 is an enzyme involved in the production of reactive oxygen species (ROS), a group of highly reactive molecules important in the regulation of signal transduction ^15^. NOX4 predominantly generates H_2_O_2_ and is constitutively active unlike the other members of the oxidase family ^16^. *NOX4* is expressed in many cell types, including keratinocytes and osteoclasts ^15^. Several studies have linked high levels of ROS to conditions such as cancer, inflammatory diseases, vascular disease, diabetes and osteoporosis ^17^. Furthermore, during bone formation, the balance between osteoblasts (bone-forming cells) and osteoclasts (bone-resorbing cells) differentiation and activity is thought to be affected by ROS ^18^. *In vitro* and *in vivo* studies have shown that increased production of NOX4 leads to increased osteoclastogenesis ^16, 19–21^. Abnormal regulation of osteoclasts activity is involved in pathological bone resorption in osteoporosis, autoimmune arthritis and bone cancer ^22^. A previous study identified an intronic SNP that increases the expression of *NOX4* being associated with reduced bone density and increased markers for bone turnover in middle-aged women compared to normal controls ^23^. Conversely, studies in mice show that depletion of NOX4 leads to increased trabecular bone density, and inhibition of NOX4 prevents bone loss ^21^. In this study, we applied massive parallel sequencing to the whole PAM cohort and found that rare variants in the *NOX4* gene found in four PAM patients might significantly increase the levels of ROS. To test the hypothesis, we applied *in vitro* and *in vivo* models, including patient-derived osteoclasts, stable cell lines over-expressing the variants found, direct measurement of superoxide in patient blood samples, and zebrafish models. Further genetic analysis of patients without *NOX4* pathogenic variants revealed other rare variants potentially pathogenic in genes related to *NOX4*. All the data obtained demonstrate a connection between higher levels of ROS and the development of PAM.

## Material and methods

### Human samples

In this study, genomic DNA was isolated from peripheral blood mononuclear cells (PBMCs) from the Nordic PAM patient’s cohort of 61 well-characterized patients previously described ^4, 6, 7, 10, 11, 24^. The cohort consists of patients from Sweden (n=27), Denmark (n=21), Norway (n=10), and Iceland (n=3). The patients’ clinical and radiographic presentations follow the consensus from the Group for Research and Assessment of Psoriasis and Psoriatic Arthritis (GRAPPA) group ^25^. In addition, for the genotyping of *NOX4* variants and for ROS measurement in blood samples we recruited psoriasis (n=1382) and psoriatic arthritis patients (n=492) and normal healthy controls (n=484). Caucasian origin was ascertained through ethnicity SNP genotyping ^26^. Blood samples from one PAM patient and one control were used for *in vitro* osteoclast differentiation.

### DNA isolation

DNA from whole blood was purified by Gentra Puregene Blood Kit (158489, Qiagen, USA). Briefly, 3 volumes RBC Lysis Solution was added to blood and centrifuged at 4000 x g for 10 minutes to pellet the white blood cells (WBS), supernatant was discarded. The WBS were lysed with 1 volume of cell lysis solution by vortexing. The cell lysates were treated with RNase A and proteins were precipitated. The supernatant containing DNA was then extracted with isopropanol followed by ethanol precipitation. After purification, the DNA was measured by Qubit.

### Whole-genome and whole exome sequence (WGS and WES)

We applied whole-genome sequencing (WGS) and whole-exome sequencing (WES) to 5 and 56 PAM patients respectively. In addition, the parents of one PAM patient were sequenced by WGS. We applied Somalier, a tool to measure relatedness in cohorts (https://github.com/brentp/somalier) to identify cryptic relatedness among all the samples (Figure S4).

WGS was performed at the Science for Life Laboratory’s (SciLifeLab) national genomics infrastructure (NGI). The sequencing libraries were constructed using 1ug of high-quality genomic DNA using the Illumina (San Diego, CA) TruSeq PCR-free kits (350 bp insert size) and sequenced on a single Illumina HiSeqX PE 2x150bp lane.

WES was conducted at Uppsala’s SNP & SEQ technological platform. We utilized 300ng of genomic DNA for WES; the DNA quality was determined using the FragmentAnalyzer, and the DNA concentration was determined using the Qubit/Quant-iT test. The sequencing libraries were constructed using the Twist Human Core Exome (Twist Bioscience), and the sequencing was carried out in a single S4 lane using the Illumina NovaSeq equipment and v1 sequencing chemicals (150 cycles paired-end).

The data were processed, and the sequence reads were aligned to the human genome build GRCh37 Single nucleotide variants (SNVs) and insertions/deletions (INDELs) were called using the GATK v3.8. and v 4.1.4.1 pipeline and the called variants were annotated using VEP (v.91). The variants were loaded into the GEMINI database to query and filter the variants. Variants with a minor allele frequency (MAF) of <0.0001 were filtered for further investigation. The variants found were inspected manually with the integrative genomics viewer (IGV) tool in the other patients. The impact of variants was evaluated using the prediction tools SIFT, Polyphen2, CADD and GERP++. Selected variants were examined manually in the BAM files using Integrated Genomics Viewer.

### Structural variants

Structural variants (SV) were analysed using FindSV, a pipeline that performs SV detection using TIDDIT and CNVnator, as well as variant filtering and annotation using VEP and SVDB ^27^. Selected variants were visualized by the Integrative Genomics Viewer (IGV) tool.

### SNP Genotyping

Genotyping of three Single Nucleotide Polymorphisms (SNPs) within the *NOX4* gene (rs781430033, rs144215891 and rs765662279) was performed by using allele-specific Taqman MGB probes labeled with fluorescent dyes FAM and VIC (Applied Biosystems, Foster City, CA, USA), according to the manufacturer’s protocols. Allelic discrimination was made with the QuantStudioTM Real-Time PCR Software (Applied Biosystems). All three mutations had to be custom-made by using Custom TaqMan®Assay Design Tool (https://www.thermofisher.com/order/custom-genomic-products/tools/cadt/); The success rate for genotyping exceeded 99% for all SNPs in the total sample set. We ran ten percent of the samples as duplicates to identify errors in genotyping and we could confirm assay accuracy of all three variations by WES and WGS of 61 PAM samples. The PCR procedure has been done using a total volume of 10ul containing 15 ng of genomic DNA, 5 ul TaqMan® Universal PCR Master Mix (2X) and 0.5 ul TaqMan® genotyping assay mix (20X). Sequences of TapMan probes and primers are listed in Table S2. Following an initial denaturation step at 50 °C for 2 min and 95°C for 10min starting all PCR procedures comprised 40 cycles of denaturation at 95 °C for 15 seconds, and primer annealing at 60 °C (55 °C for rs10065172) for 1 min and saved at 4°C. We performed an endpoint plate read comprised the last step with an increasing temperature to a maximum of 60 °C (1.6 °C per second) and accompanying measurement of fluorescence intensity on a real-time PCR on the QuantStudio 7 Flex Real-Time PCR System Instrument.

### Sanger sequencing

Genomic DNA from peripheral blood samples was extracted by standard procedures. Sanger sequencing was performed by KIGene using the ABI 3730 PRISM® DNA Analyzer ^28^. The primers used are shown in Table S1.

### Expression constructs and stable HEK293 cell lines

To test the effect of the variants in cells, we obtained plasmid - pcDNA3.1-hNox4 (#69352) from the Addgene repository. Primers with the alternate alleles for each SNP were designed using the “QuikChange Primer Design” (Agilent technologies) platform. Then, *NOX4^Y512IfsX^*^20^, *NOX4^V369F^* and *NOX4^Y512C^* variants were introduced to the construct by using QuikChange XL Site-Directed Mutagenesis Kit (Agilent) according to manufacturer instructions with the primer pairs in Table S3, which were transformed into Escherichia coli and identified by Sanger dideoxy sequencing (Table S4). To obtain stable transfectants, we linearized plasmids with 1ul BglII Enzyme (10 unit) and 5ul 10x NEB buffer with the incubation at 37°C for 15 mins and 65°C for 20 mins. HEK293 cells were kindly provided by Stefano Gastaldello (Karolinska Institutet, Stockholm, Sweden) and were transfected with pcDNA3.1-hNOX4 and three constructs carrier *NOX4* variants by the Lipofectamine 2000 Reagent (Thermo Fisher Scientific, USA), and G418 at 200 μg/ml was used as positive cell selection. Culture media containing the selection antibiotic was changed every 2-3 days until Geneticin®-resistant foci were identified. Next, we screened single-colony cells in the 96-well tissue culture plate and expanded the selected cells for future use.

### Quantitative real-time PCR analysis

The extraction of total RNA from HEK293 stable cell lines was isolated by RNeasy mini kit (QIAGEN), and cDNA was reversed with Maxima First Strand cDNA Synthesis Kit (ThermoFisher Scientific). Real-time quantitative PCR (RT-qPCR) were performed with SYBR® Green Master Mix according to the manufacturer’s protocol. Primer sequences are provided at Table S5. The 2−ΔΔCt method was utilized to achieve comparative quantification of the gene of interest between the two genotypes using actin as a reference gene.

### Western blotting

HEK293 cells were harvested and lysed in RIPA lysis with 1X Halt™ Protease and Phosphatase Inhibitor Cocktail (Thermo Fisher Scientific). The concentration of total protein was determined using the BCA Protein Assay Kit (Thermo Fisher Scientific). 10 μg or 20 μg of proteins were loaded in 10% SDS-PAGE gel and transferred to PVDF membranes. Membranes were blocked in Tris-buffered saline containing 5% skim milk for 1.5 hour at room temperature. Then incubated at 4°C overnight with recombinant anti-NADPH oxidase 4 antibody (1:2000, ab133303, abcam), and mouse anti-GAPDH monoclonal antibody was used for normalization (1:3000, 60004-1, Proteintech Group Inc). Immunoblots of protein bands were visualized with ECL (1705060, Clarity™ Western ECL Substrate, Biorad), and proteins were quantified with Image J software. The data are presented as meanJ±JSD of independent experiments performed in triplicate.

### Osteoclast studies from patient-derived peripheral blood mononuclear cells (PBMCs)

To study the effect of the mutations on osteoclasts differentiation in cell culture, we obtained patient-derived mononuclear cells. PBMCs were isolated from whole blood using Ficoll-Paque density centrifugation. For positive selection of the osteoclast precursors, i.e., the CD14+ mononuclear cells, the EasySepTM Human CD14 positive Selection kit II was used according to the manufacturer’s instructions. Purified CD14+ cells were seeded in 24-well and 96-well plates containing Gibco DMEM supplemented with 10% FBS, 0.2% PrimocinTM and macrophage colony-stimulating factor (M-CSF) (20 ng/mL; R&D systems; USA) and receptor activator of nuclear factor kappa-B ligand (RANKL) (2 ng/mL; R&D systems; USA) to induce osteoclastogenesis. Every third day, media was refreshed. The osteoclasts were fixed and stained with tartrate-resistant acid phosphatase (TRAP)-positive cells based on a leukocyte acid phosphatase kit (cat no 387A; Sigma; USA) according to the manufacturer’s instructions. TRAP-stained cells containing three or more nuclei were defined as osteoclasts ^29^.

### Measurement of superoxide by Electron Paramagnetic Resonance (EPR)

The levels of ROS in the human blood and cultured cells were measured by EPR Spectroscopy^30^. Following approximately 36 hours after transfection, cell culture media was removed, and the cells were rinsed twice with PBS. Seven hundred microliters of cyclichydroxylamine (CMH, 200 μM) in EPR-grade Krebs HEPES buffer supplemented with 25 μM Deferoxamine (DFX) and 5 μM diethyldithiocarbamate (DETC) were added to the cells and were incubated for 30 min at 37° C. The cells are collected in the 1 mL syringes and frozen in liquid nitrogen prior to measurement. To analyze ROS levels in human blood, blood samples were incubated with CMH spin probe as above mentioned and ROS was measured using the EPR spectrometer (Noxygen, Elzach, Germany). ROS levels were converted to the concentration of CP radical using the standard curve method. Briefly, blood samples were combined with cyclichydroxylamine (CMH) spin probe and ROS was measured by a CP radical standard curve, using EPR spectrometer (Noxygen, Elzach, Germany).

### Measurement of ROS production

2′,7′-dichlorofluorescein diacetate (4091-99-0, DCFH-DA, Sigma) was used as a sensitive and rapid identification of ROS in response to oxidative metabolism. Firstly, we reconstitute in DMSO for stock and then DCFH-DA S0033) was diluted with the serum-free cell culture medium. After washing osteoclasts with PBS twice at designated time points - day 8 and day 12, osteoclasts (at the density of 1×10^5^cells/well) in 96-well plate were incubated with 10 μM DCFH-DA in the incubator for 30 minutes and thereafter immediately analyzed using a fluorescence microscope (magnification ×10; EVOS^TM^ FL, Invitrogen). The relative fluorescence intensity of DCFH-DA was analyzed using Image J.

### Zebrafish assay for oxidative stress

The pcDNA3.1-h*NOX4*, *NOX4^Y512IfsX^*^20^, *NOX4^V369F^*and *NOX4^Y512C^* plasmids were linearized by restriction digestion with XhoI enzyme, and capped mRNA was transcribed *in vitro* using the mMESSAGE mMACHINE kit (Ambion, Thermo Fisher Scientific, Waltham, MA, USA). Zebrafish embryos (AB strain) at 1-2-cell stage were co-injected with *NOX4* mRNA and an antisense oligonucleotide used to knockdown endogenous nox4 expression (nox4 atg MO) (300 pg). For in vivo H2O2 detection, 30 hours post-fertilization old embryos were exposed to 20 μM 2′,7′-dichlorofluorescein diacetate (DCFH-DA, Sigma-Aldrich) for 1 hour at 28.5 °C in the dark followed by washing with embryo water a minimum of three times ^31^. ROS production was visualized by mounting each embryo in a drop of low melting agarose ^32^ and imaged using a confocal microscope (Zeiss LSM700 coupled with a water dripping lens). Importantly, every embryo was imaged using the same settings (e.g. laser intensity and optical slice thickness). The average fluorescence density (normalized to area) was analyzed using ImageJ. To this end, maximum intensity projections were produced and the total fluorescent intensity within the defined area was quantified. For each experimental group 10–16 embryos were quantified, and the experiment was repeated 3 times.

### Statistics

Quantification and Statistical Analysis were performed using GraphPad Prism 9. All experiments were performed with at least three independent biological replicates and expressed as the means ±standard deviation (SD). Student’s t-test was applied to assess the statistical differences between experimental groups. The one-way analysis of variance (ANOVA) was used to assess the statistically significant differences between the means of three unrelated groups. Multiple comparisons were evaluated for all pairs of means by Two-way ANOVA with Tukey’s correction. P☐<☐0.05 was considered significant (*p☐<☐0.05, **p☐<☐0.01, ***p☐<☐0.001, ****p☐<☐0.0001).

### Study approval

The study was approved by ethical review boards at each institution and conducted according to the Declaration of Helsinki Principles. Written informed consent was obtained from all the participants in the study. Ethical permits:2007/1088-31/4. D.nr:00-448, 2008/4:5, Dnr 02-241, and Dnr 2022-04253-02. The Stockholm Ethical Board for Animal Experiments authorized standard operating procedures for all treatments involving zebrafish (Ethical approval: Dnr 14049-2019).

## Results

### Rare *NOX4* variants in PAM patients

To investigate the pathomechanism of PAM, we looked for the presence of rare variants in patients from the Nordic PAM cohort (n=61). We applied paired-end short-read sequencing to all the patients. The parameters utilized for the study design and the filtering for rare variants are outlined in Figure 1. We found two rare variants (MAF<0.0001) in the *NOX4* gene in two Swedish patients and one variant with low frequency (MAF<0.001) in two additional patients from Denmark (Figure 2A, Table 1).

**Figure 1.**
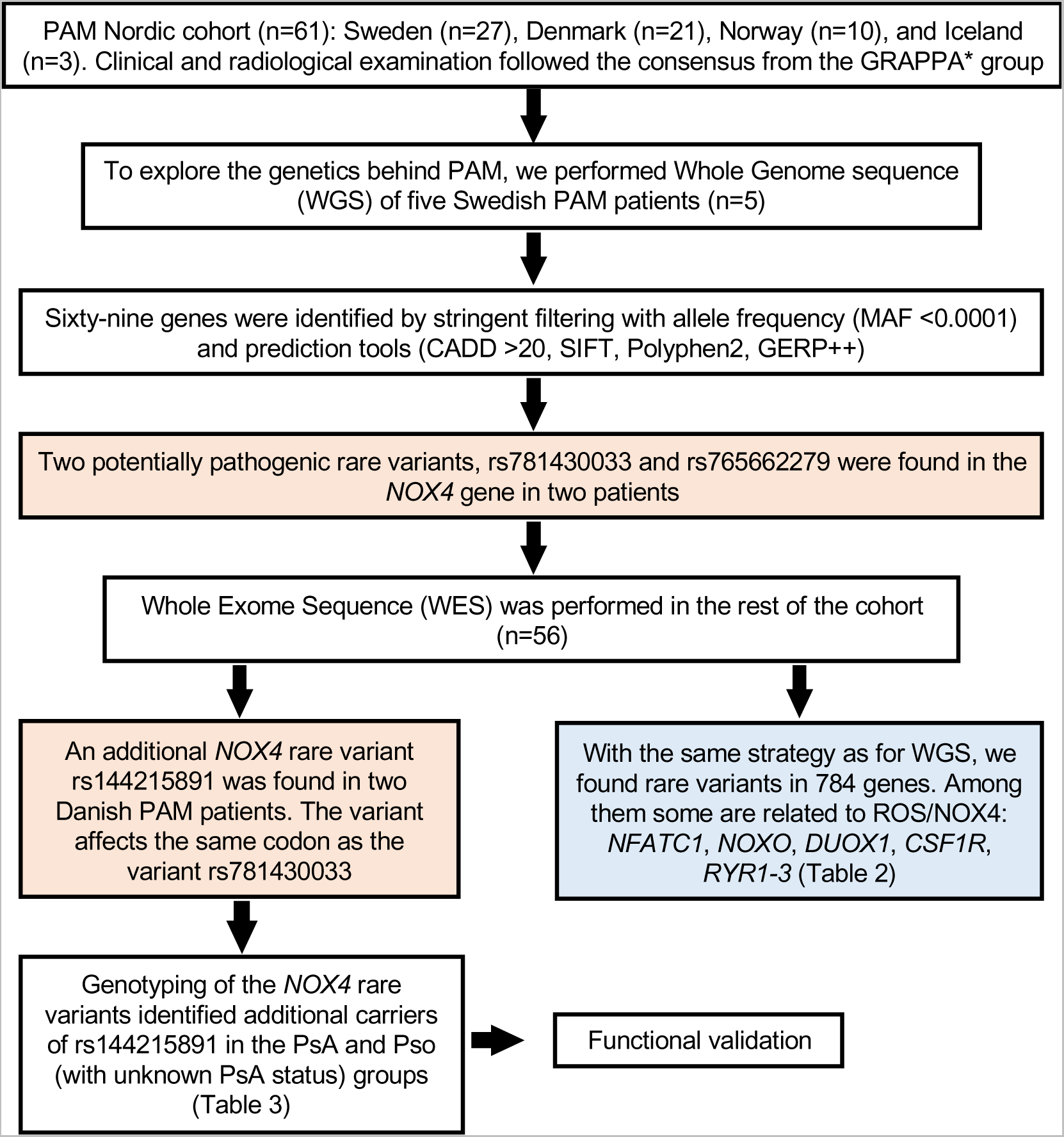
Flowchart outlining the study design and the criteria applied in the filtering of rare variants found in PAM patients by next generation sequencing. *GRAPPA: Group for Research and Assessment of Psoriasis and Psoriatic Arthritis.

**Figure 2.**
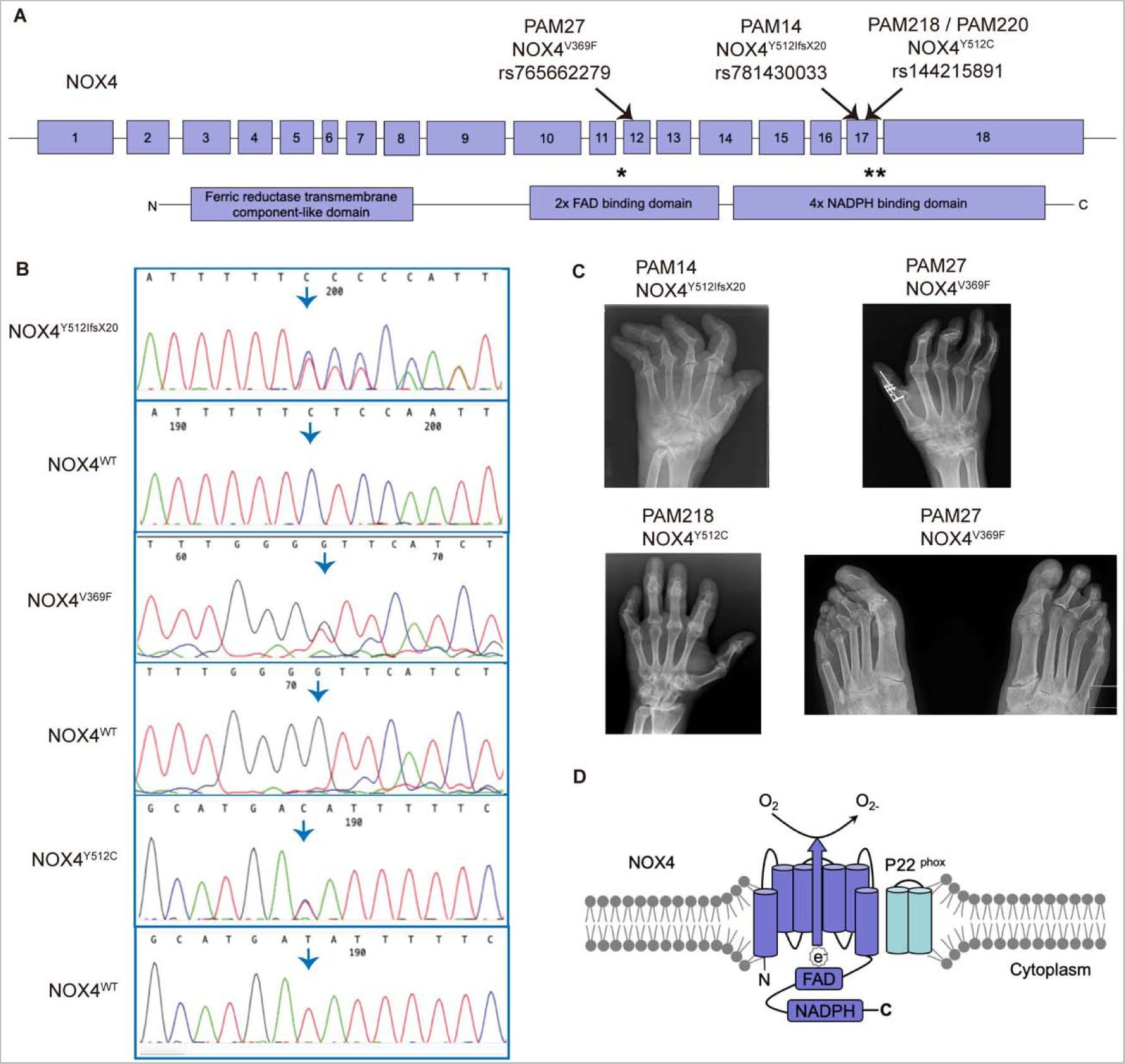
Next generation sequencing reveals rare variants in the *NOX4* gene in PAM patients. (**A**) *NOX4* gene structure is shown, exons are denoted as boxes, and the three rare variants found in PAM patients are marked by arrows; below, protein domains are indicated, stars show the position of the variants in the protein. (**B**) Sanger sequencing of the patients validated the findings from next generation sequencing, heterozygous variants in *NOX4* are shown: p.V369F, p.Y512IfsX20, and p.Y512C. Arrows highlight the nucleotide change or the start of the frameshift. (**C**) Patient radiographs of the hands show Pencil-in-cup deformities in metacarpophalangeal joints, osteolysis and ankylosis in proximal interphalangeal joints and destruction of the wrist (os carpale) (severe in PAM14, milder in PAM218). Feet: Severe osteolysis of the interphalangeal (IP) joints on the left side. Ankylosis of the first IP joint. The mutations found in patients are denoted *NOX4^Y512IfsX^*^20^, *NOX4^Y512C^*, and *NOX4^V369F^*. (**D**) Molecular structure of NOX4. Six transmembrane domains are represented by cylinders, cytosolic domains including FAD and NADPH-binding domains are shown as boxes. e, electron; FAD, flavin cofactor; NADPH, Nicotinamide Adenine Dinucleotide Phosphate.

**Table 1.**
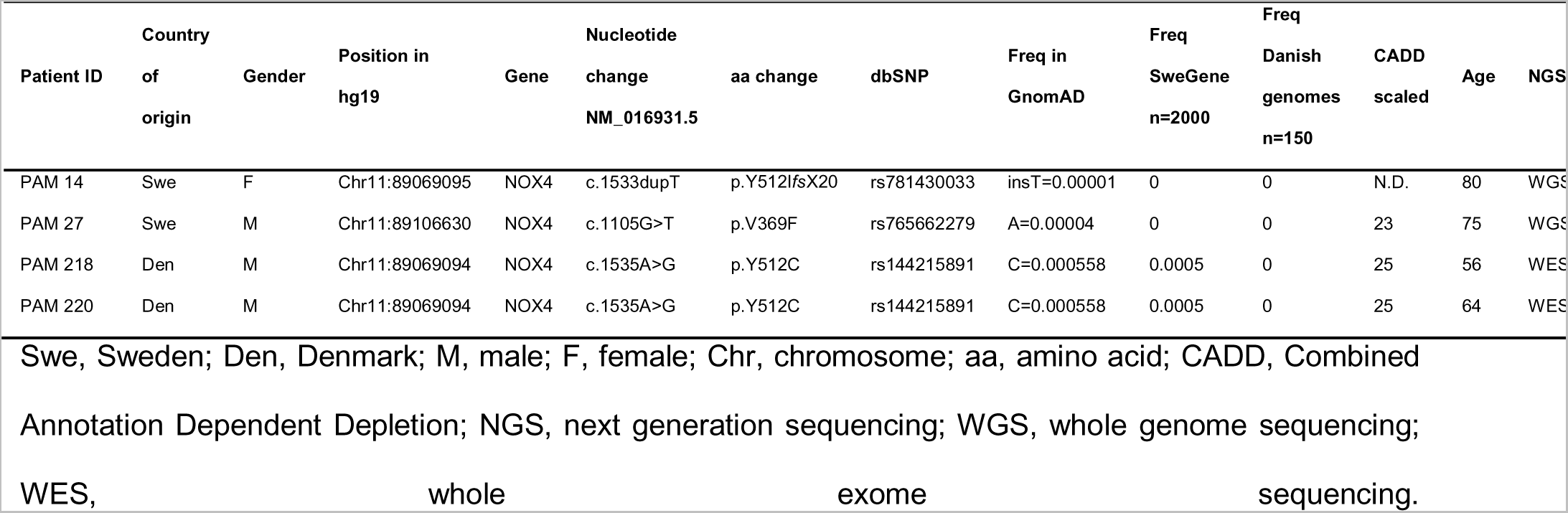
*NOX4* rare coding variants in PAM patients

One of the identified variants, rs781430033 (c.1533dup; p.Y512I*fs*X20), inserts an extra T nucleotide causing a frameshift and shortening the protein by 46 aa (Table 1). It has a frequency of 2/230742 in GnomAD and 2/120600 in ExAC databases. The second variant is a missense variant, rs765662279 (c.1105G>T; p.V369F) (Table 1), with a frequency in the population of 9/245174 in GnomAD and 4/121010 in ExAC databases. *In silico* predictions of the rs765662279 with the Combined Annotation Dependent Depletion (CADD) tool in GRCh37-v1.6 resulted in a score of 23 (a CADD score of 20 meaning that the variant is in the top 1% of most deleterious substitutions in the human genome) ^33^. The third variant - rs144215891 (c.1535A>G; p.Y512C) found in Danish patients is present at a frequency of 77/139948 in GnomAD and 71/120620 in ExAC (Table 1). The *NOX4* variants p.Y512I*fsX20* and p.Y512C are located one base pair apart from each other, and both affect the NADPH binding site region. The variant p.V369F affects the FAD binding domain (Figure 2, A and D). All variants have been confirmed by Sanger sequencing (Figure 2B).

### Radiological examination of PAM patients

Radiographical examination of three PAM patients carrying rare variants in *NOX4*, PAM14 (*NOX4^Y512IfsX^*^20^), PAM27 (*NOX4^V369F^*) and PAM218 (*NOX4 ^Y512C^*), showed typical features of PAM (Figure 2C). Patient radiographs of the hands showed shortening of the digits, severe destruction of the distal joints with pencil-in-cup deformities and osteolysis. In PAM27, severe destruction of the wrist (os carpale) was visible on the radiographs along with severe osteolysis of the interphalangeal joints in the left foot. Ankylosis of the first interphalangeal (IP) joint was also visible.

### Other rare variants related to NOX4 in PAM patients

As the *NOX4* variants were observed in limited number of patients (4 out of 61), we extended our analysis to genes involved in the ROS/NOX4 pathway. Interestingly, we found eight additional rare and potentially pathogenic variants in genes that are implicated in ROS pathways, such as *NFATC1, NOXO*, *DUOX1, CSF1R, RYR1, RYR2*, and *RYR3* summarized in Table 2.

**Table 2.**
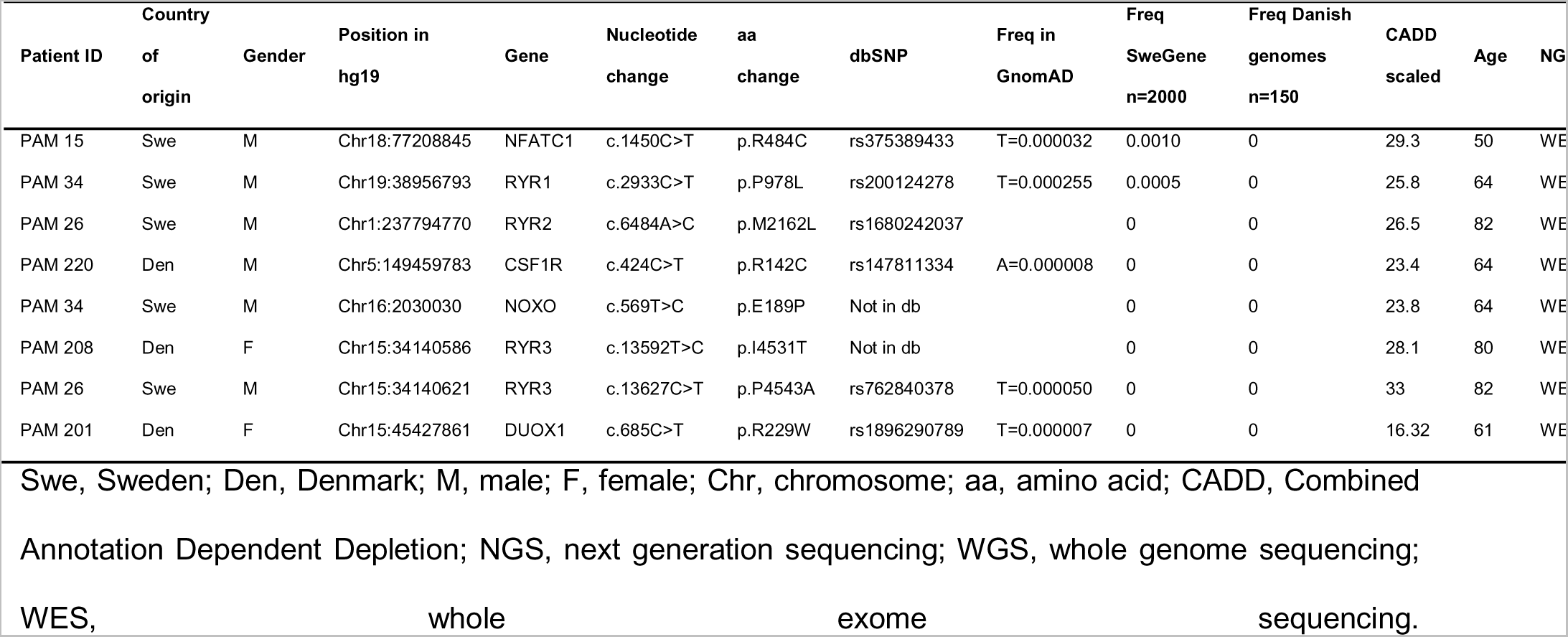
Other rare variants affecting the generation ROS in PAM patients

The nuclear transcription factor of the activated T cells c1 (*NFATC1)* gene is a key transcription factor with an essential role in osteoclast differentiation ^22^. The variant found has a high CADD score of 29.3 and the frequency in the population is very low T=0.000019 (5/264690, TOPMED) and T=0.000141 (17/120380, ExAC) (Table 2). Other potentially relevant rare variants were found in the ryanodine receptors *RYR1, RYR2*, and *RYR3*. The variants found in the *RyR* genes are extremely rare or non-existing in databases with CADD scores ranging from 25.8 to 33 and predicted to be damaging or deleterious by *in silico* tools such as PolyPhen ^34^ or SIFT ^35^. *CSF1R* is the receptor of the Colony-stimulating factor-1 (CSF-1) that is released from osteoblasts and stimulates the proliferation of osteoclast progenitors ^36^. NADPH oxidase organizer 1, *NOXO*, and Dual oxidase 1*, DUOX1* are enzymes that produce ROS ^37^ (Table 2).

We also analyzed structural variants (SVs) in the five PAM patients sequenced by whole genome sequencing. We used the FindSV pipeline to filter the variants. No SVs in *NOX4* nor in any other gene related to *NOX4* pathways were found (Supplementary File Structural Variants).

### Genotyping of *NOX4* variants in other cohorts of psoriasis, PsA and healthy controls

In order to elucidate if the variants found in *NOX4* are specific to PAM, we genotyped the three SNPs rs781430033 (NOX4^Y512IfsX^^20^), rs765662279 (NOX4^V369F^) and rs144215891 (NOX4^Y512C^) in previously described case-control cohorts of psoriasis (n=1874) and age and gender-matched healthy controls from Sweden (n=484). ^24^. We found five additional carriers of the variant NOX4^Y512C^, three in the PsA group and two in a group of 820 psoriasis patients with unknown PsA status (Table 3). In the psoriasis cases without arthritis and in healthy controls none of the variants was detected. No additional carriers of the rare variants NOX4^Y512IfsX^^20^ and NOX4^V369F^ were found in any of the groups investigated.

**Table 3.** *NOX4* rare variants in PsO, PsA and control groups

| Marker | Alleles<sup>1</sup> | PsA (N=492) | PSO No PsA<br>(N=562) | PSO unknown PsA<br>(N=820) | PAM (N=63) | Controls<br>(N=484) |
| --- | --- | --- | --- | --- | --- | --- |
| rs781430033 | A/T | 478/0/0 | 552/0/0 | 815/0/0 | 62/1/0 | 478/0/0 |
| rs765662279 | C/T | 479/0/0 | 552/0/0 | 818/0/0 | 62/1/0 | 480/0/0 |
| rs144215891 | T/C | 478/3/0 | 550/0/0 | 816/2/0 | 61/2/0 | 480/0/0 |

<sup>1</sup>Major/Minor
<sup>2</sup> PsA patients' carrying rs144215891 are: PsA961, PsA252 and PsA690

Next, in the same cohorts we investigated the *NOX4* intronic variant rs11018268 (MAF C=0.211354, GnomAD). A previous study by Goettsch et al ^23^ showed that the CC and CT genotypes of rs11018268 are associated with higher levels of *NOX4* expression, decreased bone density and an increased level of bone turnover markers ^23^. In our cohort, we found that two carriers of *NOX4^Y512C^*, patients PAM218 and PAM220, are also carriers of the rs11018268-CC and rs11018268-CT alleles respectively, and nine other PAM patients carried the CT allele. The PAM cohort was not enriched for the CC and CT alleles compared to the other groups analysed (Table S6).

### Expression of *NOX4* is higher in HEK293 cell models of *NOX4^Y512IfsX^*^20^*, NOX4^V369F^* and *NOX4^Y512C^*

To study the functional relevance of the three *NOX4* variants found in PAM, we generated HEK293 stable transfected cell lines overexpressing *NOX4* wild type (wt) and each of the three identified *NOX4* variants. The sequences of all the expression constructs were confirmed by Sanger sequencing (Figure S1). Stably transfected cells with *NOX4*^wt^ and the three rare variants *NOX4^Y512IfsX^*^20^, *NOX4^V369F^* and *NOX4^Y512C^*, were analyzed for *NOX4* expression by Real-Time qRT-PCR (Figure 3A and Figure S2). Interestingly, all three rare variants resulted in enhanced overexpression of *NOX4* mRNA compared to the overexpression of *NOX4^wt^*. The highest and most significant expression was observed for the *NOX4 ^Y512IfsX^*^20^ variant (Figure 3A). All primers used are listed in Tables S3-5.

**Figure 3.**
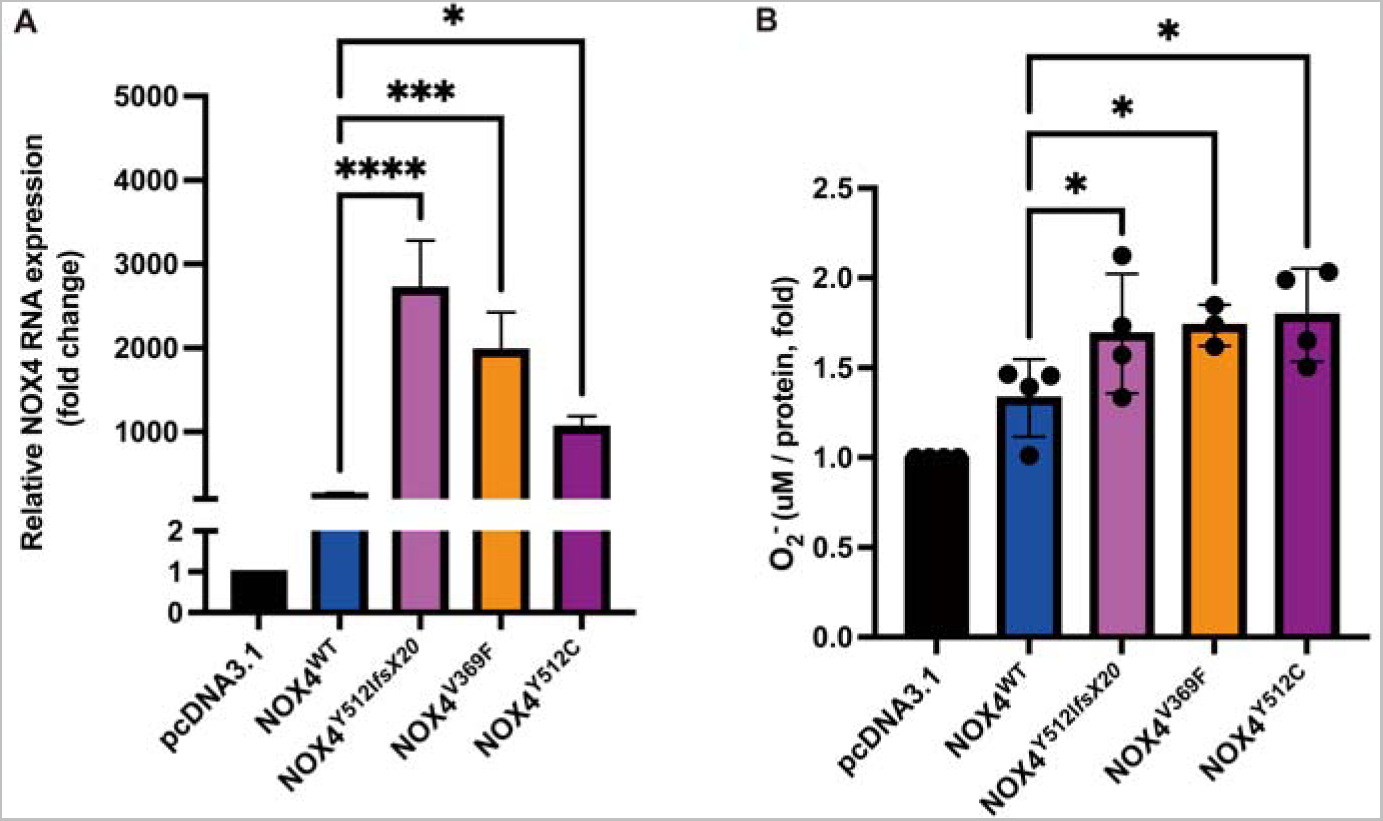
HEK293 cell lines overexpressing the rare variants found in *NOX4* increased *NOX4* expression and reactive oxygen species (ROS) generation compared to cells overexpressing *NOX4^wt^*. (**A**) The expression of *NOX4* was analyzed by qRT-PCR of cells extracts from HEK293 stable transfected cells expressing pcDNA3.1 (empty vector), *NOX4^wt^* (wild-type), *NOX4^Y512IfsX^*^20^*, NOX4^Y512C^*, and *NOX4^V369F^*. β*-actin* levels were used as a loading control. Relative levels of *NOX4* were quantified from three independent experiments and analyzed by one-way ANOVA. (**B**) All NOX4 rare variants expressing cells have higher generation of ROS compared to NOX4^wt^. Data are shown as mean ± SEM. * p < 0.05; ** p < 0.01; *** p < 0.001; **** p < 0.0001 (one-way ANOVA)

As it is known that NOX4 is involved in the ROS pathway, we hypothesized that the level of ROS might be affected. To assess the effect on ROS production in the stably transfected cell lines we performed Electron Paramagnetic Resonance (EPR). EPR is a highly sensitive and unique method that allows the direct detection of radicals ^38, 39^. We observed higher ROS levels in all cells expressing the rare variants compared to cells overexpressing *NOX4^wt^*. The increase was 1.79-fold for the *NOX4^Y512C^*, 1.69-fold for *NOX4^Y512IfsX^*^20^ and 1.74-fold for *NOX4^V369F^* compared with the empty vector (Figure 3B). Our *in vitro* results indicate that the three *NOX4* variants found in PAM patients increase the ROS levels compared to *NOX4*^wt^ in HEK293 cells.

### ROS levels are significantly higher in patients PAM12 and PsA961

To validate the involvement of the reactive oxygen species (ROS) in the development of PAM, we recruited individuals with Psoriasis (n=7), PsA (n=8), PAM (PAM12 and PAM37), as well as age and gender-matched healthy controls (n=9). ROS levels were measured in fresh peripheral blood samples by using the sensitive EPR method. ROS expression was significantly elevated in patients PAM12 and PsA961 (PsA patient carrier of *NOX4^Y512C^* detected through genotyping, see Table 3) compared to all the other groups (Figure 4A). The results suggest that the variant *NOX4^Y512C^*affecting the NADPH binding domain is responsible for the elevated superoxide production. NOX4 protein expression of PsA961 in osteoclasts is in line with this observation (Figure S3). It should be noted here that we did not find any *NOX4* mutations in PAM12 and PAM 37, nor mutations in any other genes related to *NOX4*. Analysis of PAM37 showed no significant increase in ROS production (Figure 4B). One explanation for this finding may be that patient PAM37 was treated with etanercept, an anti-TNF drug, one day before the sample collection possibly affecting the level of ROS. It should be noted here that we did not find any *NOX4* mutations in PAM12 and PAM 37, nor mutations in any other genes related to *NOX4*.

**Figure 4.**
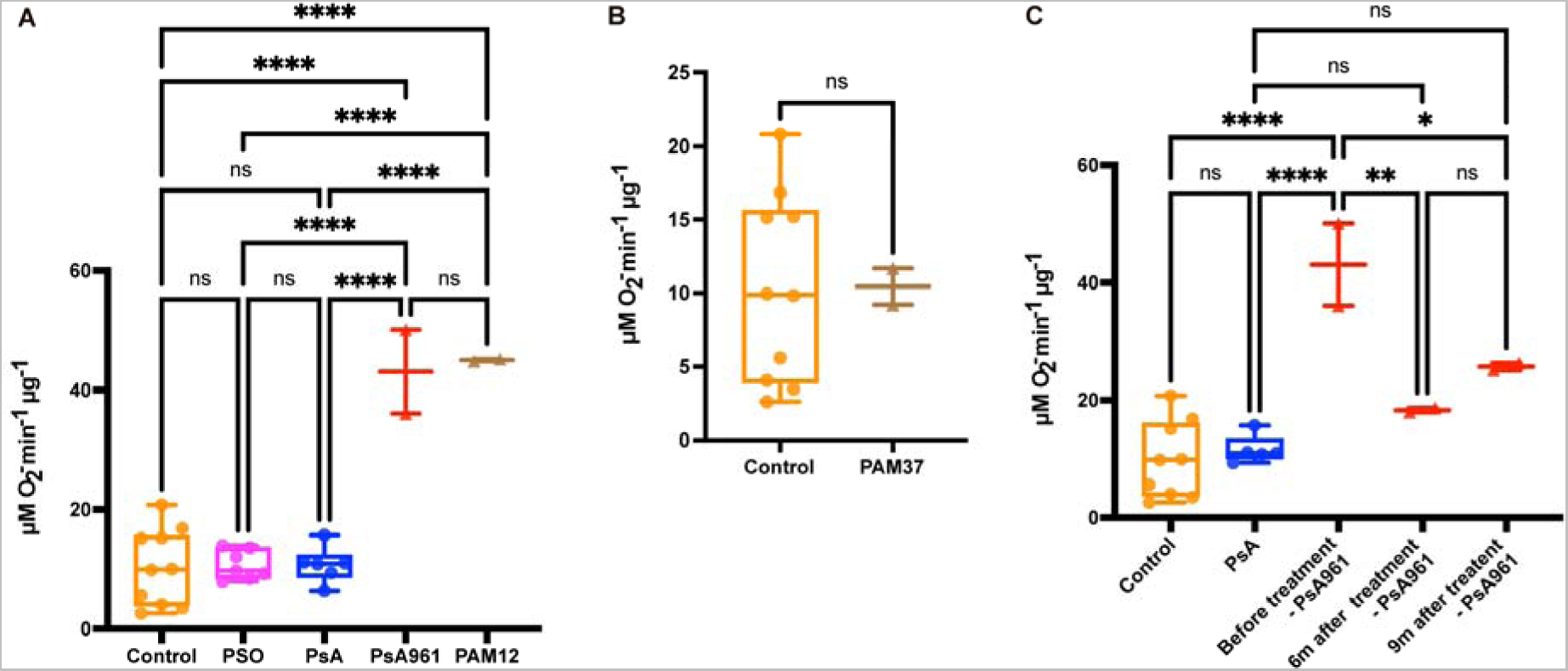
Electron paramagnetic resonance (EPR) show significant increased levels of ROS in the patients PAM12 and PsA961. (**A**) ROS measurement by EPR in peripheral blood from controls (n=9), Pso (n=7), PsA (n=5), PsA961 (*NOX4^Y512C^* carrier) and PAM12 show significant increase in PAM12 and PsA961 compared to healthy control, PSO and PsA groups. (**B**) ROS measurement of the PAM37 patient undergoing anti-TNFα treatment shows no difference compared to samples from healthy controls. In each group, circles denote samples from different individuals and in the patients PAM12 and PsA961 each triangle denotes a technical replicate. (**C**) ROS levels of the PsA961 patient decreased significantly after Adalimumab treatment. Peripheral blood samples were obtained in three occasions: before treatment, six months (6m), and nine months (9m) after treatment. * p < 0.05; ** p < 0.01; *** p < 0.001; **** p < 0.0001 (one-way ANOVA).

### The levels of ROS in patient PsA961 are decreased after treatment with adalimumab

The patient PsA961 was treated with adalimumab, a monoclonal antibody that suppresses tumor necrosis factor-alpha (TNFα) and inhibits ROS production ^40^ followed by ixekizumab for his skin psoriasis. The patient had severe skin psoriasis since 10 years and presented a very mild PsA phenotype of recent onset with intermittent pain in his wrist and knees and in one finger. Radiographs of the hands were normal at the start of adalimumab. We studied whether the overproduction of ROS in the PsA961 patient was affected by the treatment. Fresh whole blood was obtained at three different time points: before treatment started, six months, and nine months while on adalimumab treatment and 5 months following treatment start of ixekizumab. At all timepoints after biological therapy, ROS levels were normalized reaching the ROS levels of the healthy controls and other PsA patients without *NOX4* variants (Figure 4C).

### PAM12 derived osteoclasts show increased differentiation and higher generation of ROS compared to cells from a healthy control

NOX4 is induced during osteoclast differentiation, the cells responsible for bone resorption ^23^. To determine if the process of osteoclastogenesis is affected in PAM, we performed *in vitro* osteoclast differentiation of patient-derived cells from patient PAM12 and an age- and gender-matched healthy control (C12). Osteoclast differentiation was induced in CD14+ peripheral blood mononuclear cells by treatment with cytokines M-CSF and RANKL (Figure 5A). We observed a higher number of differentiated osteoclasts in the PAM12 patient compared to the healthy control, determined by the presence of multinucleated TRAP-positive cells, at both day 8 and 12 of culture (Figure 5B). To examine cellular ROS production in osteoclasts, we used 2’-7’dichlorofluorescin diacetate (DCFH-DA). DCFH-DA is a cell-permeable compound that is deacetylated by cellular esterases and oxidized by ROS into 2’-7’dichlorofluorescein (DCF). The DCF emitted fluorescence can then be quantified by fluorescence microscopy ^41^. ROS production in PAM12 osteoclasts at both days 8 and 12 of culture was significantly higher compared to C12 osteoclasts (Figure 5C). In addition, NOX4 protein levels were higher in the PAM12-derived osteoclasts compared to C12-derived osteoclasts (Figure 5D).

**Figure 5.**
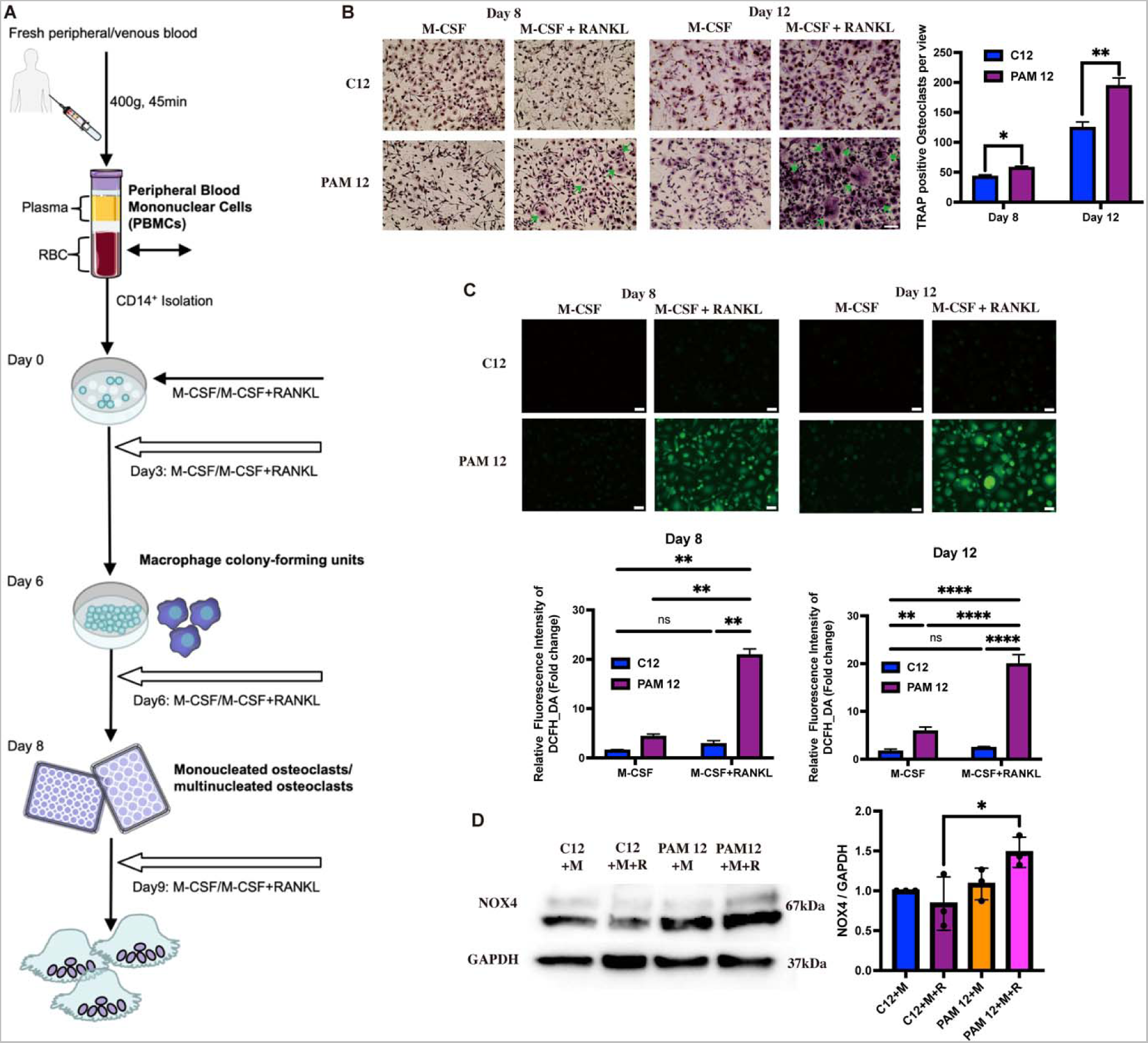
PAM12 patient derived osteoclasts show enhanced differentiation and increased ROS generation activity compared to osteoclast-derived cells from a healthy control C12. (**A**) Schematic illustration of isolation of osteoclasts from peripheral blood. More details are shown under supplementary materials and methods section. (**B**) Osteoclasts were differentiated with colony-stimulating factor (M-CSF/M) and receptor activator of nuclear factor κB ligand (RANKL/R) for 8 and 12 days. Tartrate resistant acid phosphatase (TRAP) staining (violet-labeled) was used to mark differentiated osteoclast (>3 nuclei). PAM12-derived cells show a higher number of differentiated osteoclasts (marked by arrows) compared to cells derived from a healthy control. The number of TRAP-positive osteoclasts per view was counted blindly by 2 persons. (**C**) ROS probed by DCFH-DA in cells from PAM12 patient is significantly higher compared to control cells at both 8 and 12 days after differentiation with M-CSF and RANKL. Representative images are shown. (**D**) In differentiated osteoclast (Day 8) NOX4 levels are higher in PAM12 compared to C12 control (One-way ANOVA). * p < 0.05; ** p < 0.01; *** p < 0.001; **** p < 0.0001. All the experiments were repeated at least three times (scale bars = 100 μm)

### *NOX4* rare variants *NOX4^Y512IfsX^*^20^*, NOX4^V369F^*, and *NOX4^Y512C^* increase generation of ROS in zebrafish

To further examine the effect of the three variants found in PAM patients *in vivo*, we tested the ROS production in zebrafish embryos injected with *NOX4* mRNA coding for the wild type as well as the three rare variants (Figure 6). We co-injected the mRNAs with a *nox4* translation-blocking antisense oligonucleotide, or morpholino (*nox4* atg MO), to reduce the amount of the endogenous Nox4 production and better resemble the expression patterns in patients. Protein quantification confirmed a 50% reduction in Nox4 following the injection of *nox4* atg MO (Figure 6A). Quantification of ROS by DCFH-DA revealed a higher amount of ROS production in embryos overexpressing the *NOX4* variants compared to those overexpressing *NOX4^wt^* (Figure 6C). Quantification of the fluorescence intensity in a 100 x 100 μm region of the embryos’ trunk region (Figure 6B) showed a significant increase for all three constructs NOX4^Y512IfsX^^20^, NOX4^V369F^ and NOX4^Y512C^ (Figure 6, C and D). These results in zebrafish embryos are consistent with the results obtained from HEK293T cells and patient’s cells (Figure 3A).

**Figure 6.**
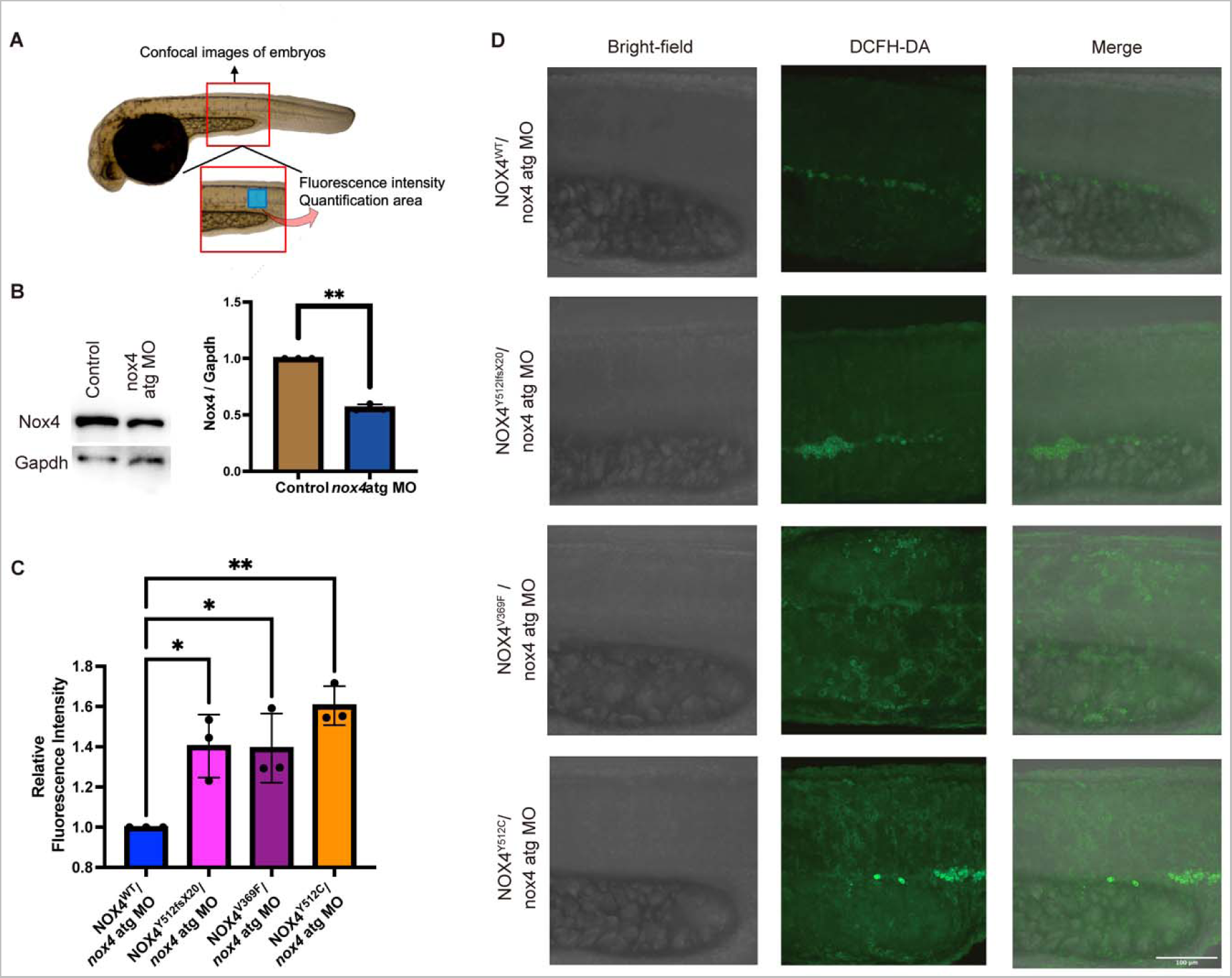
ROS production is increased in zebrafish embryos injected with *NOX4* mRNA variants compared to *NOX4^wt^*. (**A**) Endogenous Nox4 was reduced by co-injecting embryos with a translation blocking morpholino (*nox4*-atg MO), resulting in a 50% reduction of Nox4 compared to mock injected controls. Gapdh was used as loading control. Data are shown as mean ± SEM. Student’s t-test was used. (**B**) ROS production was assessed by imaging zebrafish embryos (red box) and quantifying the fluorescence intensity in the area marked by the blue box. (**C**) Quantification of fluorescence intensity show significant increase in the embryos co-injected with *NOX4* variants mRNA and nox4 atg MO compared to *NOX4*^wt^ and nox4 atg MO. Three independent experiments were measured * p < 0.05; ** p < 0.01 (Two-way ANOVA). The number of zebrafish embryos imaged in each experimental group is listed in Table S7. (**D**) Representative images of injected embryos exposed to DCFH-DA

## Discussion

The NOX gene family is comprised of seven members *NOX1-NOX5*, *DUOX1* and *DUOX2* ^42^. They are specialized ROS producers and differ in their cellular and tissue-specific distributions. Impairment in the regulation of NOX expression results in pathologies such as atherosclerosis, hypertension, diabetic nephropathy, lung fibrosis, cancers, and neurodegenerative diseases ^43^. *NOX4* specifically has been linked to osteoporosis, inflammatory arthritis and osteoarthritis ^42^. The role of *NOX4* in those pathologies has been suggested by tissue specific-expression studies, functional biochemical assays, and observations in animal models ^23, 44, 45^, but to our knowledge no disease-causing variants in *NOX4* have yet been found in patients.

Our study identified *NOX4* as the first candidate susceptibility gene for psoriatic arthritis mutilans (PAM), the rarest and most severe form of psoriatic arthritis. Despite the severity of the disease, no specific treatment nor biomarkers have been identified to date. Here, we describe three protein coding rare variants; two missense (*NOX4^V369F^* and *NOX4^Y512C^*) and one frameshift (*NOX4^Y512IfsX^*^20^), in four PAM patients, all located in the cytosolic part of NOX4, affecting the FAD and NADPH binding domains, important for the transfer of electrons and formation of ROS ^46^. In our genetic analysis, we did not find any other *NOX4* rare variants in the rest of the PAM cohort (n=57), but we cannot exclude that these patients may carry other variants located in intergenic or intragenic regions which would escape our analysis as most of the patients were sequenced only for exomes. It is also possible that other non-ROS related pathways could be implicated in the development of PAM. Through further genetic analysis of the three rare variants in psoriasis, PsA and control cohorts, we found three additional carriers of the *NOX4^Y512C^*, all in the PsA group whereas none in the psoriasis group nor in healthy controls (Table 3). Of the non-PAM patients carrying *NOX4* mutations, only PsA961 was available for further clinical examination. He presented a mild PsA phenotype affecting peripheral joints without any evidence of bone destruction. It should be noted that his PsA is of short duration, and it was his cutaneous psoriasis that motivated anti -TNF therapy. Thus, we cannot know whether he would have developed a more advanced phenotype without systemic therapeutic intervention. Also, the pathogenetic architecture in PAM is likely complex and we do not have a complete picture.

Further analysis of PAM patient sequences revealed rare variants in other genes potentially altering the levels of ROS and/or affecting osteoclast differentiation including the transcription factor *NFATc1, CSF-1R, NOXO, DUOX1, RYR1, RYR2* and *RYR3*. Interestingly, the *NFATc1* gene is a master regulator of RANKL-induced osteoclastogenesis and the *Nfatc1* conditional knockout mouse develops osteopetrosis, a condition characterized by increased bone density due to decreased or absent osteoclast activity ^47^. *CSF-1R* is involved in osteoclast proliferation, and its suppression has been shown to attenuate pathological bone resorption in inflammatory arthritis, inflammatory bone destruction, and osteoporosis ^48^. *DUOX1* is part of the NADPH family and, like *NOX4*, also produces hydrogen peroxide. Duox1 forms heterodimers with dual oxidase maturation factor 1 (Duoxa1), which was recently shown to be involved in osteoclast differentiation and ROS production in bone ^49^. Ryanodine receptors (RyRs) are calcium (Ca^2+^) channels that are responsible for Ca^2+^ release from the sarcoplasmic reticulum ^50^. In cancer-associated bone metastasis in a mouse model, upregulation of *Nox4* results in elevated oxidization of skeletal muscle proteins, including RyR1 ^51^. Also, *NOXO* is involved in ROS formation, and shown to play role in angiogenesis ^52^. Altogether, the variants affecting NOX4/ROS levels pathways are found in ∼20% of the PAM patients.

It is interesting to note that the *NOX4* intronic SNP rs11018628, previously linked to reduced bone density and elevated plasma markers for bone turnover ^23^ is found in the three PAM patients carrying NOX4 missense rare variants (*NOX4^Y512C^, NOX4^V369F^*), but not in the PAM patient carrying the frameshift variant (*NOX4^Y512IfsX^*^20^). We also found the SNP rs11018628 in the PsA961 carrier of *NOX4^Y512C^*. Perhaps, there could be an additive or synergistic effect of these variants on *NOX4* expression at the transcriptional level, leading to increased generation of ROS. Our analysis of the ROS levels in patient PsA961 using electron paramagnetic resonance (EPR) showed significantly increased ROS levels compared to other individuals in PsA, psoriasis and healthy controls groups (Figure 4A). The ROS levels were similar to the levels observed in the PAM12 patient. Unfortunately, we were not able to recruit any of the other patients for the measurement.

Several factors are involved in the regulation of *NOX4*, including NF-κB, TGF-β, TNFα, endoplasmic reticulum (ER) stress, hypoxia, and ischemia but the underlying mechanisms behind the regulation are not fully understood ^53^. Our results are in line with the previous observations showing that upregulation of NOX4 is linked to several pathogenic conditions, such as idiopathic pulmonary fibrosis ^54^, chronic obstructive pulmonary disease ^55^ several cardiovascular conditions ^56^ and osteoporosis ^23^.

In summary, we here present novel genetic findings, supported by several lines of functional evidence for the involvement of ROS in the etiology of PAM: *i)* using stably transfected HEK293 cells, we show that the rare variants result in elevated *NOX4* transcript expression and ROS generation (Figure 3, A and B), *ii)* measurement of ROS in patient PAM12 (a patient without identified *NOX4* mutations) and patient PsA961 (carrier of *NOX4^Y512C^*) showed a significant increase of ROS compared to control, psoriasis and PsA samples (Figure 4, A and B), *iii)* patient-derived cells from PAM12 showed increased osteoclast differentiation with increased ROS activity compared to cells from a healthy control (Figures 5, B and C), and finally *iv)* using a zebrafish model, we show *in vivo* that the generation of ROS is significantly enhanced by all three *NOX4* rare variants found in PAM patients (Figures 6, C and D).

A limitation of the present study is the lack of access to fresh blood samples from the PAM patients, needed for measurement of ROS by EPR. Our study would have benefited from deeper exploring the ROS levels in more patients. Nevertheless, we had the possibility of testing a couple of PAM patients and a PsA (carrier of *NOX4^Y512C^*) by EPR as a proof of concept that the generation of ROS is indeed affected in both patients. Another consideration is that the rare variants found in *NOX4* are observed in just a few PAM patients (4 out of 61). Additional genetic analysis indicates potentially pathogenic variants in other genes found in PAM patients also affecting osteoclast differentiation and activity. Further functional validation experiments are required to test the pathogenicity of those variants.

Interestingly, the patients at risk of developing PAM may benefit from existing biological therapies applied in moderate and severe psoriasis which may reduce the generation of ROS. Another commonly used drug for treating psoriasis, methotrexate, inhibits osteoclast differentiation by inhibiting RANKL ^57^. With the advent of effective therapies for psoriasis and psoriatic arthritis, PAM has become increasingly rare, still it is important to early diagnose and treat to avoid irreversible damage.

This study reveals a direct link to NOX4 and ROS production in PAM pathology and gives the first strong indication of where to search for specific disease identifiers in this destructive disease. Would early intervention with existing biologic treatments be sufficient or is precision therapy essential? The disease process can be rapid in PAM resulting in irreparable damage. Early identification of those at risk and initiation of effective therapy would constitute a game changer.

## Supplemental information

Supplemental data

Supplementary file Structural Variants

## Supporting information

Supplemental structural variants

## Acknowledgments

We gratefully acknowledge the patients and controls for participation in this project. We also would like to thank Helena Griehsel for helping in taking samples from patients; Jose Laffita-Mesa, Anton Tornqvist and Xiaoyuan Ren for technical assistance and Andrea Bieder for providing critical editorial feedback on the manuscript.

The authors acknowledge support from the National Genomics Infrastructure in Stockholm funded by Science for Life Laboratory, the Knut and Alice Wallenberg Foundation and the Swedish Research Council, and SNIC/Uppsala Multidisciplinary Center for Advanced Computational Science for assistance with massively parallel sequencing and access to the UPPMAX computational infrastructure. We also acknowledge the support provided by the Biomedicum Imaging Core and Zebrafish Core Facility employees in maintaining the microscopes and caring for the zebrafish.

This work was supported by Hudfonden (grants 3378, 3227 and 2808 to ITP, MS and to PN), Swedish Rheumatism Association, Reumatikerförbundet (R-968063 to ITP), Konung Gustaf V:s 80-årsfond (FAI-2021-0819a to ITP), Psoriasisfonden to M.S. and ITP, The European academy of dermatology and venereology (EADV) (PPRC-2022-40 to ITP), Doctoral scholarship KI-China scholarship Council (CSC) programme to SW, Stiftelsen Sällsyntafonden to SW, FT, and RV.

## Author contributions

MS and ITP conceived and designed the study. SW, ITP, PN, RV, FT, XZ, and JJA developed and optimized the experimental part of the methodology. FT, and JE assisted with the bioinformatic analysis. SW and RV performed and analyzed the zebrafish experiments. SW, and XZ performed the EPR measurements. PN performed the genotyping. LL, BG, LE, LI, UL, SBC, and MS, provided clinical resources for the project. LL, and UL, assisted with clinical data analysis. MS, ITP, PN, and LI acquired the funding. ITP, MS, and PN performed the project management. ITP, SW, and MS wrote the original draft. All authors reviewed and approved the final manuscript.

## Data availability

All data supporting the conclusions in the article are presented in the main text or supplementary data files. Additional data are available from the corresponding author upon request.

