## Supplemental data for "Rare coding variants in *NOX4* link high superoxide levels to psoriatic arthritis mutilans"

| SNP | Forward | Reverse |
| --- | --- | --- |
| rs781430033/ rs144215891 | CAAAAGTTTCCACCGAGGACG | ACAGCAATTTGGTGGGAAGCC |
| rs765662279 | AATTATTACATTCCACTAT | GTTGTAATTGAATTATAATT |

**Table S1. Primers used for genomic DNA Sanger sequencing.**

**Table S2. Primers and probes used for Genotyping.**

| SNP | Genotyping PCR primers | Genotyping probes |
| --- | --- | --- |
| rs781430033 | F: CGAGGACGTCCTATAAACAGTCTTG  R: GTAGTGCTTTCTTTTTGGCAGAAGAT | VIC reporter: CATGATATTTTTTCTCC  FAM reporter: GCATGATATTTTTCTCC |
| rs144215891 | F: CGAGGACGTCCTATAAACAGTCTTG  R: GTAGTGCTTTCTTTTTGGCAGAAGAT | VIC reporter: AGTGCATGATATTTTTC  FAM reporter: TGCATGACATTTTTC |
| rs765662279 | F: CTACATACCTGTCCAGTCTCCTACT  R: GTGTCCAACTGAAACCAAAGCA | VIC reporter: AAGATGAACCCCAAATGT  FAM reporter: AAGATGAAACCCAAATGT |

F: forward; R: reverse

**Table S3. Primers used for site-directed mutagenesis.**

| Plasmids | Mutagenesis primers |
| --- | --- |
| NOX4**^Y512IfsX20^** | oligonucleotide primer #1: GAATTCAGTGCATGATATTTTTTCTCCAATTATCTTCTGTATCCCATCT  oligonucleotide primer #2: AGATGGGATACAGAAGATAATTGGAGAAAAAATATCATGCACTGAATTC |
| NOX4**^V369F^** | oligonucleotide primer #1: GTCTCCTACTATTTTAAGATGAAACCCAAATGTTGCTTTGGTTTCAG  oligonucleotide primer #2: CTGAAACCAAAGCAACATTTGGTTTCATCTTAAAATAGTAGGAGAC |
| NOX4**^Y512C^** | oligonucleotide primer #1: TGAATTCAGTGCATGACATTTTTCTCCAATTATCTTCTGTATCCCATC  oligonucleotide primer #2: GATGGGATACAGAAGATAATTGGAGAAAAATGTCATGCACTGAATTCA |

**Table S4. Primers used for Sanger sequencing of plasmids**

| Target gene | Primer sequence (5’-3’) | |
| --- | --- | --- |
|  | Forward | Reverse |
| NOX4**^Y512IfsX20^** / NOX4**^Y512C^** | CTTCCGTTGGTTTGCAGATT | TGGGTCCACAACAGAAAACA |
| NOX4**^V369F^** | TCCCTCAGATGTCATGGAAATC | TGAAGGGCAGAATTTCGGAG |

**Table S5. Primers used for qPCR**

| Target gene | Primer sequence (5’-3’) | |
| --- | --- | --- |
|  | Forward | Reverse |
| ACTB | CAACCGCGAGAAGATGAC | AGGAAGGCTGGAAGAGTG |
| NOX4_exon1_3 | CTGTGTCCTGGAGGAGCTGG | AAGCCAAGAGTGTTCGGCAC |
| NOX4_exon15_18 | CTTCCGTTGGTTTGCAGATT | TGGGTCCACAACAGAAAACA |

**Table S6. Genotyping of the *NOX4* variant- rs11018268 in PsO, PsA and control groups.**

| Alleles^1^ | Controls (N=451) | PsO No PsA (N=562) | PsA (N=492) | PAM (N=63) |
| --- | --- | --- | --- | --- |
| T/C | 364/75/1 | 462/87/3 | 400/74/4 | 52/10/1 |
| ^1^Major/Minor | | | | |

**Table S7.** **Number of zebrafish embryos imaged in each experimental group.**

| Number | NOX4**^WT^**/  *nox4* atg MO | NOX4**^Y512fsX20^**/  *nox4* atg MO | NOX4**^V369F^**/  *nox4* atg MO | NOX4**^Y512C^**/  *nox4* atg MO |
| --- | --- | --- | --- | --- |
| First experiment | 11 | 10 | 11 | 16 |
| Second experiment | 11 | 13 | 10 | 12 |
| Third experiment | 11 | 10 | 11 | 9 |

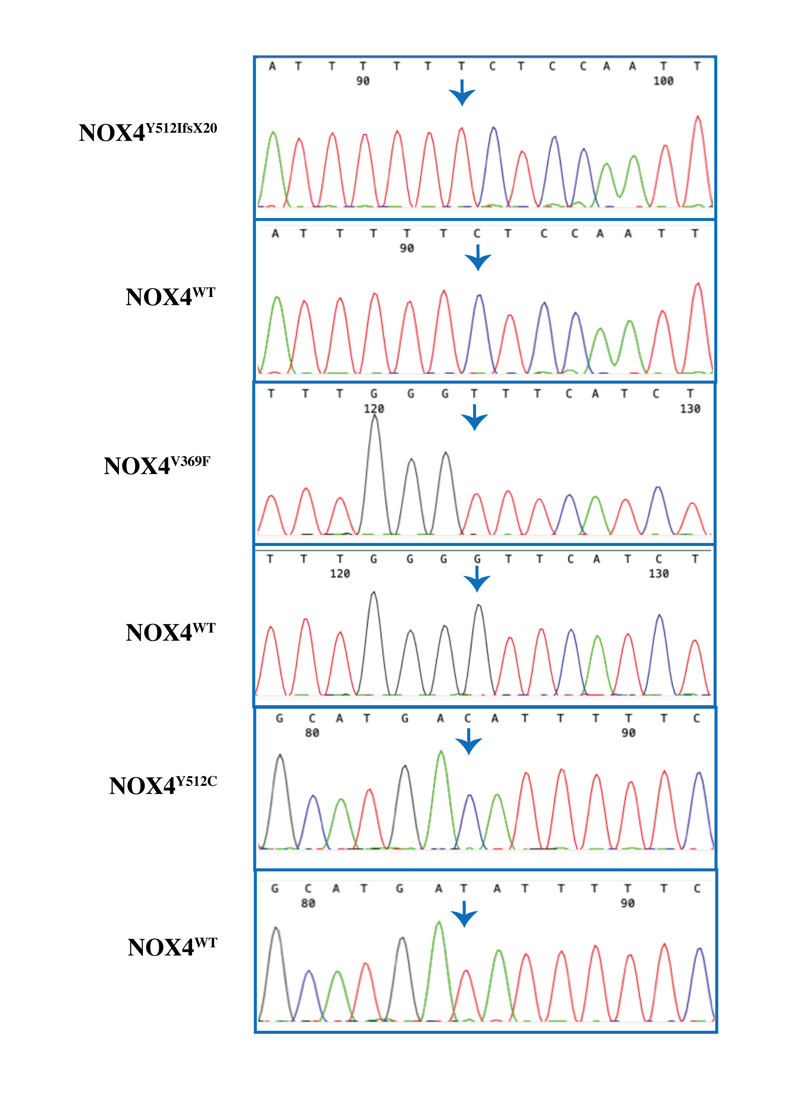

**Figure S1. Sanger sequencing results from plasmid constructs**. Forward-sequencing result of the plasmid NOX4^Y512IfsX20^ showing an extra T nucleotide insertion, which caused a frameshift. Forward-sequencing result of the plasmid NOX4^V369F^ showing Phenylalanine (F) replaced by Valine (V) on protein. Forward-sequencing result of the plasmid NOX4^Y512C^ showing Cystine (C) replaced by Tyrosine (Y) on protein. The arrow indicates the vector/insert junction. DNA bases are color-coded: A, green; G, black; C, blue; and T, red.

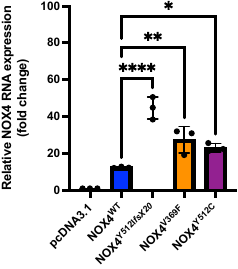

**Figure S2. NOX4 mRNA expression in HEK293 stable cells.** Primers were designed between exons 1-3. Quantitative real-time PCR analysis of the NOX4 from wild-type or NOX4-variants expressing HEK293 cells. The quantification of stable cells with NOX4 mutants were up-regulated compared with WT. Relative levels were normalized to β-actin (one-way ANOVA). Data were obtained from three independent experiments. * p < 0.05, ** p < 0.01; **** p < 0.0001.

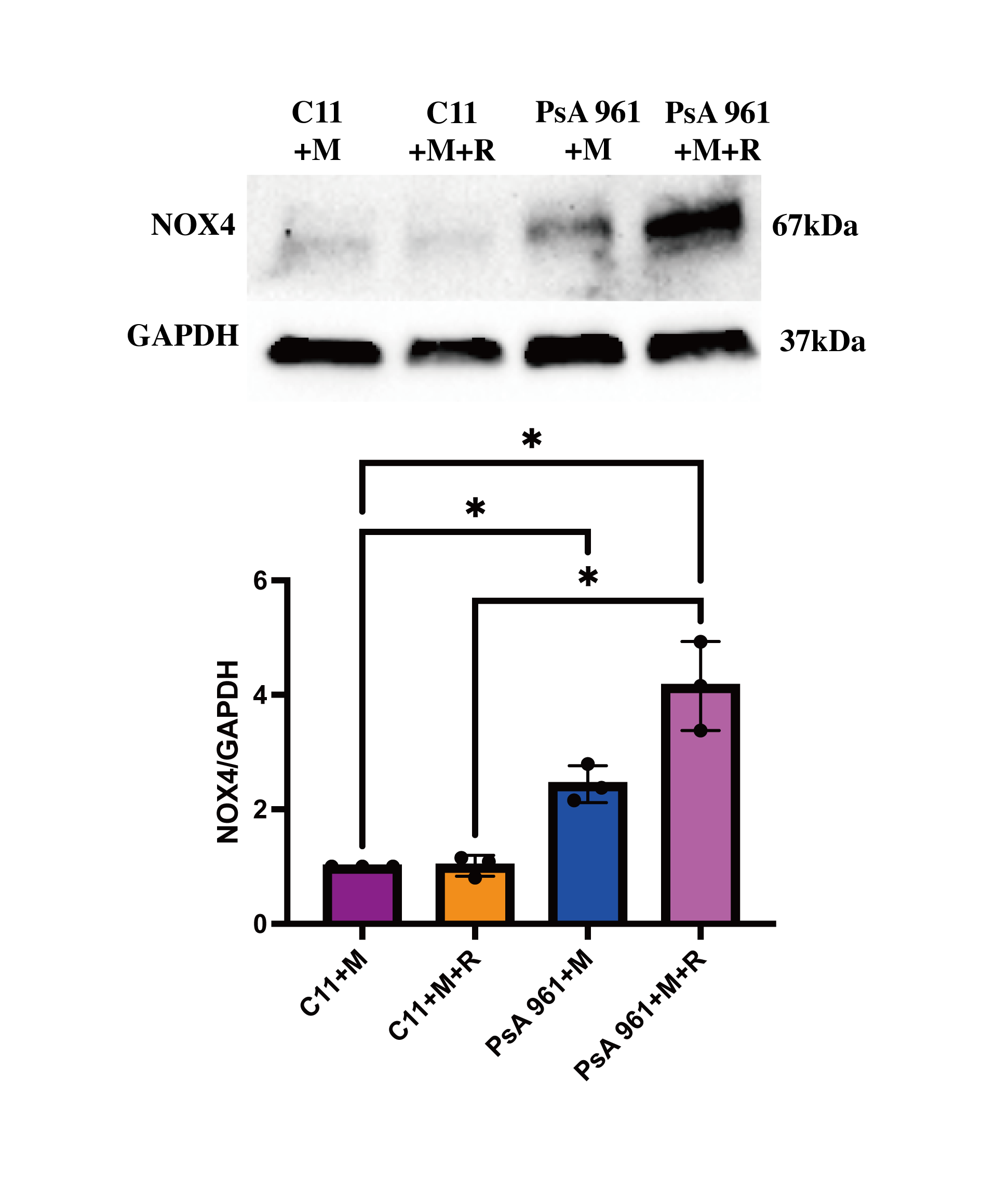

**Figure S3. The NOX4 protein expression of osteoclasts in PsA961.** Osteoclasts in PsA 961 were cultured induced from mononuclear cells with M-CSF and RANKL. Western blot imagines for expression of NOX4 were analyzed based on the results of by using an ECL detection system. NOX4 protein expression was increased after inducing by M-CSF and RANKL. GADPH levels were used as a loading control (n=3). * p < 0.05. Two-way ANOVA with Tukey’s correction for multiple testing is used.

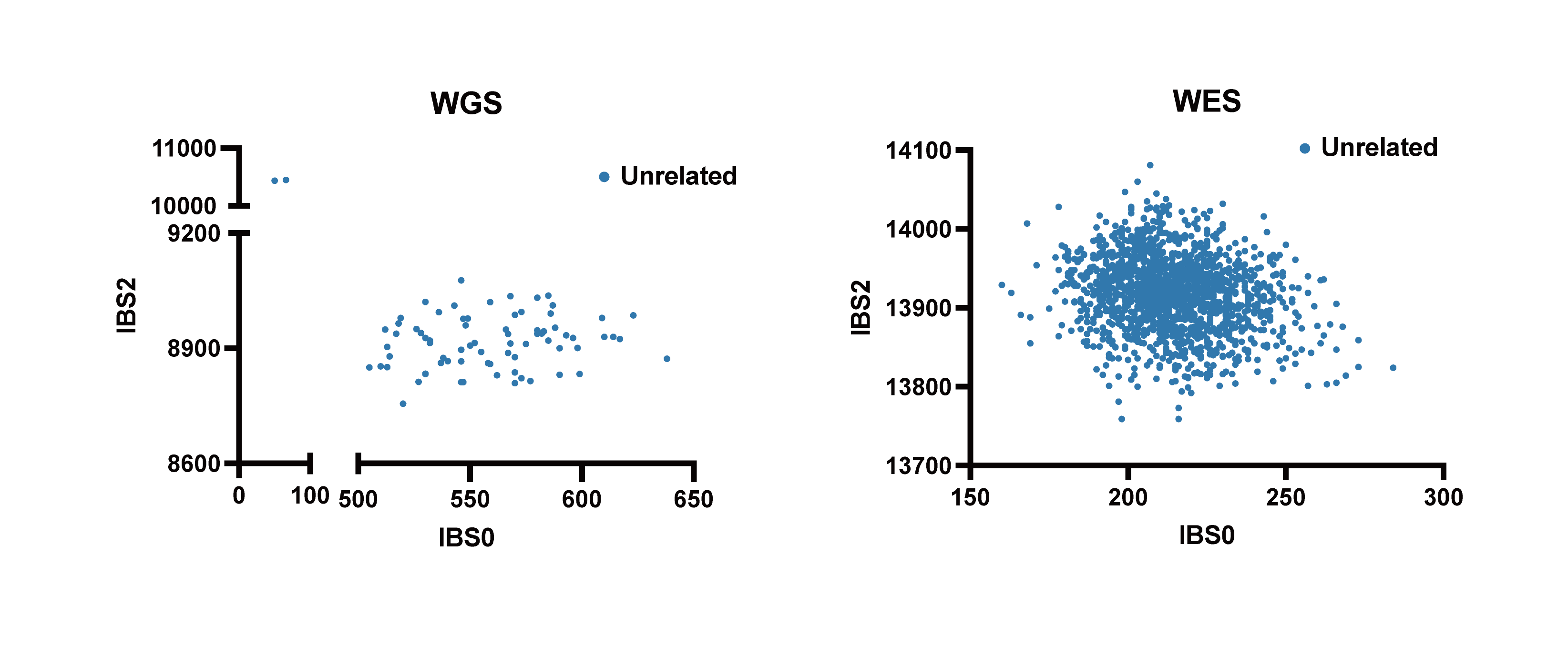

**Figure S4. Relatedness plot for WGS (A) and WES (B) samples**. Each dot represents a pair of samples. In A the samples that are in the top left are a trio (PAM12 and its parents), six samples from the SweGene database were added to the WGS analysis.  IBS0 is the number of sites where 1 sample is homozygous for the reference allele and the other is homozygous for the alternate allele. IBS2, is the count of sites where a pair of samples were both homozygous or both heterozygous. B, analysis of all WES samples.
