## Supplemental structural variants for "Rare coding variants in *NOX4* link high superoxide levels to psoriatic arthritis mutilans"

| ChromosomeA | chromosomeB | PosA | PosB | Length | variant | P10051_101 | Genes |
|---|---|---|---|---|---|---|---|
| 1 | 1 | 151001 | 156000 | 4999 | DEL | Het | RNU6-1100P\|RP11-34P13.9\|RP11-34P13.13 |
| 1 | 1 | 724823 | 224200107 | 223475284 | BND | Het | RP11-206L10.9 |
| 1 | 1 | 12994001 | 13006000 | 11999 | DEL | Het | PRAMEF6 |
| 1 | 1 | 25643001 | 25650000 | 6999 | DEL | Het | C1orf63\|RHD |
| 1 | 1 | 28736601 | 28736727 | 126 | BND | Het | PHACTR4 |
| 1 | 1 | 144308001 | 144313000 | 4999 | DEL | Het | RNVU1-4\|LINC00623 |
| 1 | 1 | 145092950 | 145097082 | 4132 | BND | Het | SEC22B\|RP4-725K1.1 |
| 1 | 1 | 206516001 | 206534000 | 17999 | DUP | Het | SRGAP2 |
| 1 | 1 | 232160608 | 232168295 | 7687 | DUP | Het | DISC1 |
| 2 | 2 | 110702001 | 110735000 | 32999 | DEL | Het | AC013271.3 |
| 2 | 2 | 131251001 | 131261000 | 9999 | DEL | Het | RNU6-473P\|POTEI |
| 2 | 2 | 131615001 | 131621000 | 5999 | DEL | Het | ARHGEF4 |
| 2 | 2 | 132519001 | 132524000 | 4999 | DEL | Het | C2orf27A |
| 2 | 2 | 132977791 | 132980445 | 2654 | BND | Het | ANKRD30BL |
| 2 | 8 | 231446295 | 5640097 | #NUM! | BND | Het | AC010149.4 |
| 3 | 3 | 125461001 | 125467000 | 5999 | DEL | Het | OR7E97P |
| 15 | 3 | 48761438 | 136910706 | #NUM! | BND | Het | \|FBN1 |
| 3 | 3 | 175062880 | 175062986 | 106 | DUP | Het | NAALADL2 |
| 3 | 3 | 195429512 | 195429622 | 110 | BND | Het | MIR570\|LINC00969 |
| 4 | 4 | 10544914 | 10552747 | 7833 | DEL | Het | CLNK |
| 4 | 4 | 33846043 | 33846339 | 296 | BND | Het | RP11-79E3.3 |
| 4 | 4 | 140775822 | 140775952 | 130 | BND | Het | MAML3 |
| 5 | 7 | 41029254 | 93840084 | #NUM! | BND | Het | MROH2B |
| 5 | 5 | 67983768 | 123167789 | 55184021 | BND | Hom | |
| 19 | 6 | 21905704 | 21766294 | #NUM! | BND | Het | VN1R84P\|ZNF100\|CASC15 |
| 6 | 6 | 45184954 | 45185101 | 147 | BND | Het | SUPT3H |
| 6 | 6 | 57284913 | 57289357 | 4444 | BND | Het | PRIM2 |
| 6 | 6 | 57423088 | 57429120 | 6032 | BND | Het | PRIM2 |
| 7 | 7 | 15463159 | 15463295 | 136 | BND | Het | AGMO |
| 7 | 7 | 44663097 | 44663213 | 116 | BND | Het | OGDH |
| 7 | 7 | 64593001 | 64598000 | 4999 | DEL | Het | INTS4L1 |
| 7 | 7 | 74958001 | 74978000 | 19999 | DUP | Het | Y_RNA\|AC006014.10\|PMS2P2\|AC004878.3 |
| 7 | 7 | 143967001 | 144024000 | 56999 | DEL | Het | CTAGE8\|OR2A9P\|ARHGEF35\|ARHGEF34P\|OR2A1\|RP4-545C24.1\|OR2A1-AS1 |
| 8 | 8 | 12251001 | 12258000 | 6999 | DEL | Het | DEFB109P1\|FAM66A |
| 9 | 9 | 5036576 | 5105455 | 68879 | BND | Het | RP11-39K24.2\|JAK2 |
| 9 | 9 | 5403904 | 5404264 | 360 | DEL | Het | PLGRKT |
| 9 | 9 | 38814001 | 38820000 | 5999 | DEL | Het | RP11-402N8.1 |

| | | | | | | | |
|---|---|---|---|---|---|---|---|
| 9 | 9 | 39185001 | 39534000 | 348999 | DEL | Het | RP11-347J14.7\|RP11-381G8.1\|FAM74A5\|CTD-2173L22.4\|RN7SL640P\|CNTNAP3\|RP11-347J14.4\|SPATA31A1\|RP11-347J14.8 |
| 9 | 9 | 39539001 | 39550000 | 10999 | DEL | Het | |
| 9 | 9 | 40703001 | 40708000 | 4999 | DEL | Het | RP11-395E19.2\|SPATA31A3\|RP11-395E19.5 |
| 9 | 9 | 41037001 | 41042000 | 4999 | DEL | Het | CTD-2340F8.3 |
| 9 | 9 | 41321001 | 41329000 | 7999 | DEL | Het | RP11-95K23.3\|SPATA31A4 |
| 9 | 9 | 43046001 | 43075000 | 28999 | DUP | Het | SNX18P4\|RP11-327I22.1\|ANKRD20A3 |
| 9 | 9 | 44145001 | 44151000 | 5999 | DEL | Het | MEP1AP4 |
| 9 | 9 | 44847001 | 44852000 | 4999 | DEL | Het | Y_RNA |
| 9 | 9 | 45132001 | 45202000 | 69999 | DEL | Het | RP11-449H15.2 |
| 9 | 9 | 45581001 | 45599000 | 17999 | DEL | Het | |
| 9 | 9 | 66768001 | 66790000 | 21999 | DUP | WT | |
| 9 | 9 | 67092001 | 67103000 | 10999 | DEL | Het | AL591379.1 |
| 9 | 9 | 99064832 | 99065206 | 374 | DEL | Het | HSD17B3 |
| 10 | 10 | 46933001 | 46939000 | 5999 | DEL | Het | FAM35BP\|RP11-38L15.2 |
| 10 | 10 | 77111864 | 77112096 | 232 | BND | Het | RP11-399K21.11\|ZNF503-AS1 |
| 10 | 10 | 134165820 | 134165941 | 121 | DUP | Het | LRRC27 |
| 11 | 11 | 3118148 | 3118271 | 123 | DUP | Hom | OSBPL5 |
| 11 | 11 | 48367530 | 48373694 | 6164 | BND | Het | OR4C4P |
| 11 | 11 | 114433315 | 131230464 | 16797149 | BND | Het | NXPE1 |
| 11 | 11 | 120321480 | 120321637 | 157 | BND | Het | ARHGEF12 |
| 12 | 12 | 11245001 | 11251000 | 5999 | DEL | Het | TAS2R14\|TAS2R43\|PRR4\|RP11-673D15.7 |
| 12 | 12 | 13545001 | 13552000 | 6999 | DEL | Het | C12orf36 |
| 12 | 12 | 54147589 | 54147701 | 112 | BND | Het | RP11-686F15.2\|RP11-686F15.3 |
| 14 | 14 | 69780350 | 69780484 | 134 | BND | Het | GALNT16 |
| 15 | 15 | 20950001 | 20988000 | 37999 | DUP | Het | RP11-403B2.3\|RP11-403B2.6\|RP11-403B2.7\|RP11-403B2.5 |
| 15 | 15 | 83024001 | 83039000 | 14999 | DEL | Het | RPL9P8\|UBE2Q2P3 |
| 16 | 16 | 18415001 | 18420000 | 4999 | DEL | Het | NPIPA8 |
| 16 | 16 | 28459001 | 28464000 | 4999 | DEL | Het | snoU13\|NPIPB7\|RP11-57A19.5 |
| 16 | 16 | 28698001 | 28703000 | 4999 | DEL | Het | EIF3C\|RP11-57A19.4 |
| 16 | 16 | 32753001 | 32769000 | 15999 | DUP | Het | HERC2P5\|RP11-586K12.5\|RP11-586K12.4 |
| 16 | 16 | 33126001 | 33165000 | 38999 | DUP | Het | FAM108A10P\|HERC2P8\|RP11-19N8.6\|RP11-1437A8.2\|RP11-19N8.7 |
| 16 | 16 | 33853087 | 33853203 | 116 | DUP | Hom | RP11-598D12.2 |
| 16 | X | 47666773 | 11731509 | #NUM! | BND | Het | PHKB |
| 16 | 16 | 77747522 | 77747752 | 230 | BND | Het | AC092724.1 |
| 17 | 17 | 20408001 | 20416000 | 7999 | DEL | Het | KRT16P3\|AC025627.9\|RP11-434D2.7 |
| 17 | 17 | 21321279 | 21322820 | 1541 | BND | Het | KCNJ12 |
| 17 | 17 | 21349691 | 25331283 | 3981592 | BND | Het | PDLIM1P2 |
| 17 | 17 | 35954262 | 35954364 | 102 | BND | Het | SYNRG |
| 17 | 17 | 72729211 | 72729477 | 266 | BND | Het | RAB37 |

| | | | | | | | |
|---|---|---|---|---|---|---|---|
| 18 | 18 | 50488067 | 50489494 | 1427 | BND | Het | DCC |
| 18 | 18 | 54493289 | 54493508 | 219 | BND | Het | WDR7 |
| 19 | 19 | 5761476 | 5761631 | 155 | BND | Het | CATSPERD |
| 19 | 19 | 6995404 | 6995509 | 105 | DUP | Het | EMR4P |
| 20 | 20 | 135121 | 136537 | 1416 | BND | Het | DEFB127 |
| 20 | 20 | 26209794 | 26210981 | 1187 | DUP | Hom | MIR663A |
| 21 | 21 | 11002001 | 11008000 | 5999 | DEL | Het | TPTE\|BAGE2 |
| 21 | 21 | 14817744 | 14869837 | 52093 | DUP | Het | GTF2IP2\|VN1R8P |
| 22 | 22 | 20418001 | 20423000 | 4999 | DEL | Het | PI4KAP1 |
| 22 | 22 | 20451001 | 20456000 | 4999 | DEL | Het | RN7SKP131\|SCARNA18\|SCARNA17\|RIMBP3 |
| X | X | 301001 | 604000 | 302999 | DUP | WT | SHOX\|PPP2R3B\|AL732314.1\|KRT18P53\|FABP5P13 |
| X | X | 606001 | 1035000 | 428999 | DUP | WT | SHOX\|RP11-309M23.1\|RPL14P5 |
| X | X | 103150001 | 103155000 | 4999 | DEL | Hom | RP11-370B6.1 |
| X | X | 145891001 | 145896000 | 4999 | DEL | Het | CXorf51B\|CXorf51A |
| X | X | 148634001 | 148640000 | 5999 | DEL | Het | CXorf40A\|RP5-937E21.8 |
| Y | Y | 13256001 | 13280000 | 23999 | DUP | WT | RP1-85D24.1\|RP1-85D24.2\|RP1-85D24.3 |
| MT | MT | 1 | 16568 | 16567 | BND | Hom | MT-ND1\|MT-ND2\|MT-TL1\|MT-TQ\|MT-TV\|MT-RNR1\|MT-RNR2\|MT-TM\|MT-TI\|MT-TF |

Structural Variants

| ChromosomeA | chromosomeB | PosA | PosB | Length | variant | P10051_102 | P10051_106 | P10051_107 | Genes |
|---|---|---|---|---|---|---|---|---|---|
| 1 | 1 | 150001 | 155000 | 4999 | DEL | Het | Het | WT | RNU6-1100P\|RP11-34P13.13 |
| 1 | 1 | 1,04E+08 | 1,04E+08 | 12999 | DUP | Het | WT | WT | AMY1C |
| 1 | 1 | 1,44E+08 | 1,44E+08 | 4999 | DEL | Het | WT | Het | LINC00623 |
| 1 | 1 | 1,45E+08 | 1,45E+08 | 4132 | BND | Het | Het | Het | SEC22B\|RP4-725K1.1 |
| 1 | 1 | 1,82E+08 | 1,82E+08 | 123 | BND | Het | WT | WT | RGSL1 |
| 10 | 10 | 2895053 | 2913524 | 18471 | DUP | Het | WT | Het | |
| 10 | 10 | 38938001 | 38994000 | 55999 | DUP | Het | WT | WT | ACTR3BP5\|SLC9B1P3 |
| 10 | 10 | 46740001 | 46748000 | 7999 | DEL | Het | WT | WT | BMS1P1\|RNA5SP311 |
| 10 | 10 | 46751001 | 46762000 | 10999 | DEL | Het | WT | WT | BMS1P1 |
| 10 | 10 | 46763001 | 46775000 | 11999 | DEL | Het | WT | WT | BMS1P1\|GLUD1P7\|DUSP8P3 |
| 10 | 10 | 47724001 | 47729000 | 4999 | DEL | Het | Het | WT | CTGLF11P |
| 10 | 10 | 49000001 | 49010000 | 9999 | DEL | Het | WT | WT | RP11-508M1.7 |
| 10 | 10 | 54287637 | 78377068 | 24089431 | BND | Het | WT | WT | |
| 10 | 10 | 67392195 | 67778937 | 386742 | DEL | Het | WT | Het | AC022538.1\|RP11-222A11.1\|CTNNA3 |
| 11 | 11 | 13816371 | 13842537 | 26166 | BND | Het | Het | WT | RP11-98J9.1 |
| 11 | 11 | 48367530 | 48373694 | 6164 | BND | Het | Het | Het | OR4C4P |
| 11 | 11 | 54804001 | 54822000 | 17999 | DUP | Het | WT | WT | |
| 11 | 11 | 1,14E+08 | 1,31E+08 | 16797152 | BND | Het | Het | Het | NXPE1 |
| 12 | 12 | 34690001 | 34719000 | 28999 | DUP | Het | WT | WT | |
| 12 | 12 | 50271294 | 50271405 | 111 | BND | Het | Het | WT | FAIM2 |
| 12 | MT | 51432311 | 7465 | #NUM! | BND | Het | WT | Het | RNU6-1273P |
| 12 | 12 | 1,14E+08 | 1,14E+08 | 165 | BND | Het | Het | WT | DTX1 |
| 14 | 14 | 72310001 | 72316000 | 5999 | DEL | Het | WT | WT | RP6-114E22.1 |
| 15 | 15 | 24502001 | 24508000 | 5999 | DEL | Het | WT | WT | RP11-580I1.2 |
| 15 | 15 | 28832001 | 28852000 | 19999 | DUP | Het | WT | WT | HERC2P9 |
| 15 | 3 | 34511629 | 1,66E+08 | #NUM! | BND | Het | WT | WT | |
| 16 | 16 | 78001 | 83000 | 4999 | DEL | Het | WT | Het | IL9RP3\|Z84812.4\|WASIR2 |
| 16 | 16 | 33076001 | 33184000 | 107999 | DUP | Het | WT | WT | FAM108A10P\|RP11-19N8.3\|RP11-19N8.4\|RP11-19N8.6\|RP11-19N8.7\|HERC2P8\|RP11-1437A8.2 |
| 16 | 16 | 69953036 | 69953143 | 107 | BND | Het | WT | WT | WWP2 |
| 16 | 16 | 80328307 | 80390590 | 62283 | DEL | Het | WT | Het | RP11-525K10.3 |
| 17 | 17 | 20353001 | 20359000 | 5999 | DEL | Het | WT | WT | LGALS9B\|NOS2P3 |
| 17 | 17 | 21321279 | 21322821 | 1542 | BND | Het | Het | Het | KCNJ12 |
| 17 | 17 | 70698004 | 70698153 | 149 | BND | Het | WT | WT | SLC39A11 |
| 17 | 17 | 78711001 | 78717000 | 5999 | DEL | Het | Het | WT | RPTOR |

| | | | | | | | | | |
|---|---|---|---|---|---|---|---|---|---|
| 17 | 17 | 78711076 | 78718605 | 7529 | BND | Het | Het | WT | RPTOR |
| 17 | 17 | 79399740 | 79399873 | 133 | BND | Hom | WT | Hom | RP11-1055B8.7 |
| 18 | 18 | 50488067 | 50489494 | 1427 | BND | Het | Het | Het | DCC |
| 19 | 19 | 125001 | 156000 | 30999 | DEL | Het | WT | WT | AC016626.1\|AC016626.2\|OR4F8P |
| 19 | 19 | 3556866 | 3556991 | 125 | BND | Het | WT | Het | AC005786.7\|AC005786.5\|MFSD12 |
| 19 | 19 | 17646077 | 17647735 | 1658 | DEL | Het | WT | Het | FAM129C |
| 19 | 19 | 18552971 | 18553096 | 125 | DUP | Hom | WT | WT | AC010335.1\|CTD-3137H5.1\|ISYNA1\|ELL |
| 2 | 2 | 71564777 | 71564881 | 104 | BND | Het | WT | WT | ZNF638 |
| 2 | 2 | 86950001 | 86956000 | 5999 | DEL | Het | WT | WT | CHMP3\|RMND5A\|RNF103-CHMP3 |
| 2 | 9 | 1,25E+08 | 93647000 | #NUM! | BND | Het | WT | WT | |
| 2 | 2 | 2,16E+08 | 2,16E+08 | 104 | BND | Het | WT | WT | AC072062.1 |
| 2 | 3 | 2,42E+08 | 1,72E+08 | #NUM! | BND | Het | WT | WT | KIF1A |
| 20 | 20 | 135121 | 136537 | 1416 | BND | Het | Het | Het | DEFB127 |
| 20 | 20 | 26209794 | 26210981 | 1187 | DUP | Hom | WT | WT | MIR663A |
| 20 | 20 | 62822669 | 62827260 | 4591 | DEL | Het | Het | WT | MYT1 |
| 22 | 22 | 20465001 | 20472000 | 6999 | DEL | Het | WT | WT | XXbac-B33L19.10\|RIMBP3 |
| 22 | 22 | 45677910 | 45678081 | 171 | BND | Het | WT | WT | UPK3A |
| 3 | 3 | 1,16E+08 | 1,16E+08 | 48413 | DEL | Het | Het | WT | TUSC7\|LSAMP |
| 4 | 4 | 7512270 | 7512451 | 181 | DUP | Het | WT | WT | SORCS2 |
| 4 | 4 | 79341642 | 79341743 | 101 | BND | Het | Het | WT | FRAS1 |
| 4 | 4 | 1,36E+08 | 1,36E+08 | 4999 | DEL | Het | WT | WT | RP11-553P9.2 |
| 5 | 5 | 36145305 | 36146296 | 991 | BND | Het | WT | Het | LMBRD2\|MIR580 |
| 5 | 5 | 68835001 | 68840000 | 4999 | DEL | Het | WT | WT | OCLN\|snoU13\|GUSBP3 |
| 5 | 5 | 68950001 | 68986000 | 35999 | DEL | Het | WT | WT | RP11-974F13.5\|GUSBP3 |
| 5 | 5 | 69473001 | 69479000 | 5999 | DEL | Het | WT | WT | RP11-1415C14.4 |
| 5 | 5 | 1,73E+08 | 1,73E+08 | 40595 | DUP | Het | WT | Het | CTB-43E15.3\|AC008674.1 |
| 5 | 5 | 1,75E+08 | 1,75E+08 | 112 | DUP | Hom | WT | Hom | RP11-826N14.1 |
| 5 | 5 | 1,77E+08 | 1,77E+08 | 4999 | DEL | Het | WT | WT | FAM153A |
| 5 | 5 | 1,8E+08 | 1,8E+08 | 175 | BND | Het | WT | WT | CNOT6 |
| 5 | 5 | 1,81E+08 | 1,81E+08 | 11999 | DEL | Het | WT | WT | |
| 6 | 6 | 3145774 | 3146181 | 407 | BND | Het | WT | WT | RP1-40E16.11\|BPHL |
| 6 | 6 | 57284913 | 57289357 | 4444 | BND | Het | Het | Het | PRIM2 |
| 6 | 6 | 57423088 | 57429121 | 6033 | BND | Het | Het | Het | PRIM2 |
| 6 | 6 | 1,71E+08 | 1,71E+08 | 79999 | DEL | Het | WT | Het | XX-C2158C12.1\|OR4F7P |
| 7 | 7 | 15463159 | 15463295 | 136 | BND | Het | WT | WT | AGMO |
| 7 | 7 | 27783417 | 27783724 | 307 | DUP | Hom | WT | WT | TAX1BP1\|AC004549.6 |
| 7 | 7 | 44663097 | 44663213 | 116 | BND | Het | Het | WT | OGDH |

| | | | | | | | | | |
|---|---|---|---|---|---|---|---|---|---|
| 7 | 7 | 65436617 | 65465712 | 29095 | DUP | Het | Het | WT | RP11-252P18.1\|GUSB |
| 7 | 7 | 74716001 | 74875000 | 158999 | DEL | Het | WT | WT | AC004878.7\|AC138783.12\|AC004878.2\|Y_RNA\|AC004878.8\|GATSL2\|AC118138.2 |
| 7 | 7 | 1,01E+08 | 1,01E+08 | 1066 | DEL | Het | WT | WT | MUC17 |
| 7 | 7 | 1,31E+08 | 1,31E+08 | 174 | BND | Het | WT | WT | MKLN1 |
| 7 | 7 | 1,38E+08 | 1,38E+08 | 220 | DEL | Het | Het | WT | ATP6V0A4 |
| 7 | 7 | 1,42E+08 | 1,42E+08 | 8999 | DEL | Het | WT | WT | MGAM |
| 8 | 8 | 7089001 | 7100000 | 10999 | DEL | Het | WT | WT | AF228730.13\|OR7E125P |
| 8 | 8 | 13958887 | 13959074 | 187 | BND | Het | WT | WT | SGCZ |
| 8 | X | 17740802 | 1,03E+08 | #NUM! | BND | Het | WT | WT | RP11-156K13.2\|FGL1 |
| 8 | 8 | 18451656 | 18451851 | 195 | BND | Het | WT | WT | PSD3 |
| 8 | 8 | 43132272 | 43133719 | 1447 | DEL | Het | WT | Het | RP11-726G23.10\|RP11-726G23.7 |
| 8 | 8 | 1,36E+08 | 1,36E+08 | 116 | BND | Het | WT | WT | RP11-1057B8.2 |
| 8 | 8 | 1,42E+08 | 1,42E+08 | 124 | BND | Het | WT | WT | SLC45A4 |
| 9 | 9 | 78001 | 86000 | 7999 | DEL | Het | WT | WT | RP11-143M1.2\|RP11-143M1.7 |
| 9 | 9 | 43871001 | 43877000 | 5999 | DEL | Het | WT | WT | CNTNAP3B |
| 9 | 9 | 43903001 | 43910000 | 6999 | DEL | Het | WT | WT | CNTNAP3B |
| 9 | 9 | 44431001 | 44740000 | 308999 | DEL | Het | WT | WT | ATP5A1P6\|RP11-475I24.6\|AL162415.4\|AL162415.2\|AL162415.1\|RBPJP6 |
| 9 | 9 | 44847001 | 44852000 | 4999 | DEL | Het | WT | WT | Y_RNA |
| 9 | 9 | 66344001 | 66350000 | 5999 | DEL | Het | WT | WT | CNN2P5\|RP11-459O16.1 |
| GL000209.1 | GL000209.1 | 39001 | 70000 | 30999 | DEL | Het | WT | WT | KIR2DL2 |
| MT | MT | 1 | 16568 | 16567 | BND | Hom | Hom | Hom | MT-ND1\|MT-ND2\|MT-TL1\|MT-TQ\|MT-TV\|MT-RNR1\|MT-RNR2\|MT-TM\|MT-TI\|MT-TF |
| X | X | 1892063 | 1892261 | 198 | DUP | Het | WT | WT | RP13-297E16.5 |
| X | X | 2548142 | 2548324 | 182 | INV | Het | WT | WT | CD99P1 |
| X | X | 3751001 | 3760000 | 8999 | DEL | Het | WT | WT | RP11-706O15.1 |
| X | X | 52886897 | 52887007 | 110 | BND | Het | WT | WT | XAGE3 |
| X | X | 1,35E+08 | 1,35E+08 | 196815 | BND | Hom | Het | WT | RP11-432N13.2\|DDX26B |
| X | X | 1,35E+08 | 1,35E+08 | 90457 | INV | Het | Het | WT | RP11-432N13.3\|RP11-432N13.2\|DDX26B\|RP11-432N13.4 |

Structural Variants

| ChromosomeA | chromosomeB | PosA | PosB | Length | variant | P10051_103 | Genes |
|---|---|---|---|---|---|---|---|
| 1 | 1 | 13156001 | 13161000 | 4999 | DEL | Het | RP13-221M14.3 |
| 1 | 1 | 13646001 | 13653000 | 6999 | DEL | Het | PRAMEF15 |
| 1 | 1 | 13655001 | 13665000 | 9999 | DEL | Het | PRAMEF14 |
| 1 | 1 | 145092950 | 145097082 | 4132 | BND | Het | SEC22B\|RP4-725K1.1 |
| 1 | 1 | 159045251 | 159045364 | 113 | DUP | Het | AIM2 |
| 1 | 1 | 161562001 | 161598000 | 35999 | DUP | Het | HSPA7\|FCGR3A\|FCGR2B\|FCGR2C\|RPS23P9\|FCGR3B\|RP11-25K21.6 |
| 1 | 1 | 219923452 | 219923572 | 120 | DEL | Het | SLC30A10 |
| 2 | 2 | 88174001 | 88186000 | 11999 | DUP | Het | RGPD2 |
| 2 | 2 | 111011001 | 111018000 | 6999 | DEL | Het | RP11-1223D19.1 |
| 2 | 2 | 131228001 | 131234000 | 5999 | DEL | Het | POTEI |
| 3 | 3 | 8601110 | 8601815 | 705 | BND | Het | LMCD1\|LMCD1-AS1 |
| 4 | 4 | 25334143 | 25334767 | 624 | DEL | Het | ZCCHC4 |
| 4 | 9 | 38735213 | 96386715 | #NUM! | BND | Het | |
| 4 | 4 | 170542628 | 170542779 | 151 | BND | Het | CLCN3 |
| 4 | 4 | 178657744 | 178657848 | 104 | BND | Het | LINC01099\|LINC01098 |
| 4 | 4 | 190945001 | 190951000 | 5999 | DEL | Het | FRG2\|DUX4L9 |
| 5 | 5 | 17207101 | 17208655 | 1554 | DEL | Het | AC091878.1\|DCAF13P2\|BASP1 |
| 5 | 5 | 106231958 | 106232058 | 100 | BND | Het | CTC-254B4.1 |
| 5 | 5 | 175471253 | 175471365 | 112 | DUP | Hom | RP11-826N14.1 |
| 5 | 5 | 177154001 | 177160000 | 5999 | DEL | Het | FAM153A |
| 6 | 6 | 57284913 | 57289357 | 4444 | BND | Het | PRIM2 |
| 6 | 6 | 57423088 | 57429121 | 6033 | BND | Het | PRIM2 |
| 6 | 6 | 57573001 | 57598000 | 24999 | DUP | WT | |
| 6 | 6 | 65218724 | 65219390 | 666 | DEL | Het | EYS |
| 6 | 6 | 149895297 | 149895448 | 151 | BND | Het | RP1-12G14.6\|GINM1 |
| 7 | 7 | 57014001 | 57019000 | 4999 | DEL | Het | RP11-220H4.2\|MIR4283-2 |
| 7 | 7 | 74375001 | 74380000 | 4999 | DEL | Het | GATSL1 |
| 8 | 8 | 13958887 | 13959074 | 187 | BND | Het | SGCZ |
| 8 | 8 | 43434001 | 43453000 | 18999 | DUP | Het | |
| 9 | 9 | 42163001 | 42286000 | 122999 | DEL | Het | RP11-216M21.5\|RP11-216M21.4\|RP11-216M21.1\|RP11-216M21.2 |
| 10 | 10 | 2896001 | 2990000 | 93999 | DEL | Het | RP11-89K18.1 |
| 10 | 10 | 2896049 | 3137672 | 241623 | INV | Het | RP11-118K6.2\|RP11-89K18.1\|RP11-118K6.3\|PFKP |

| | | | | | | | |
|---|---|---|---|---|---|---|---|
| 10 | 10 | 2989918 | 3112666 | 122748 | INV | Het | RP11-118K6.2\|RP11-118K6.3\|PFKP |
| 10 | 10 | 3112001 | 3138000 | 25999 | DUP | Het | RP11-118K6.3\|PFKP |
| 10 | 10 | 42936703 | 43021729 | 85026 | INV | Het | LINC00839\|ZNF37BP\|CCNYL2 |
| 10 | 10 | 47913001 | 48301000 | 387999 | DEL | Het | GLUD1P6\|CTSLP2\|SLC9A3P4\|DUSP8P2\|ANXA8\|ASAH2C\|FAM21B\|AL591684.1\|AGAP9\|RP11-301J7.9\|RNA5SP313\|FAM25G\|RP11-301J7.8\|BMS1P6 |
| 11 | 11 | 48367530 | 48373694 | 6164 | BND | Het | OR4C4P |
| 11 | 11 | 114433315 | 131230467 | 16797152 | BND | Het | NXPE1 |
| 12 | 12 | 50271294 | 50271405 | 111 | BND | Het | FAIM2 |
| 12 | 12 | 50292024 | 50292132 | 108 | BND | Het | FAIM2 |
| 13 | 13 | 19067001 | 19122000 | 54999 | DUP | Het | BNIP3P7\|LINC00349 |
| 14 | 14 | 80511644 | 80538313 | 26669 | DEL | Het | |
| 15 | 15 | 22344001 | 22511000 | 166999 | DUP | Het | IGHV1OR15-3\|IGHV1OR15-1\|MIR1268A\|snoU13\|IGHV1OR15-4\|IGHV4OR15-8\|RP11-69H14.6\|RP11-2F9.4\|OR4N3P\|OR4N4\|RP11-603B24.6\|OR4H6P\|RP11-2F9.3\|AC010760.1\|OR4M2 |
| 15 | 15 | 22513001 | 22596000 | 82999 | DUP | Het | RP11-603B24.1\|RP11-603B24.2\|MIR1268A |
| 15 | 15 | 28860001 | 28897000 | 36999 | DUP | Het | HERC2P9 |
| 15 | 15 | 32463001 | 32468000 | 4999 | DEL | Het | Y_RNA\|CHRNA7 |
| 15 | 15 | 32817001 | 32823000 | 5999 | DEL | Het | RP11-632K20.7 |
| 15 | 15 | 32873001 | 32879000 | 5999 | DEL | Het | RP11-1000B6.3 |
| 16 | 16 | 16706001 | 16795000 | 88999 | DEL | Het | RP11-14N9.2\|RN7SL90P |
| 16 | 16 | 18651001 | 18757000 | 105999 | DUP | Het | |
| 16 | 16 | 28458001 | 28464000 | 5999 | DEL | Het | snoU13\|NPIPB7\|RP11-57A19.5 |
| 16 | 16 | 32217001 | 32658000 | 440999 | DUP | WT | ABCD1P3\|RP11-56L13.1\|RP11-586K12.11\|RP11-17M15.2\|RP11-17M15.4\|RP11-626K17.5\|RP11-626K17.3\|RP11-626K17.2\|PABPC1P13\|RP11-96K14.1\|RP11-56L13.6\|RP11-586K12.13\|TP53TG3D\|ACTR3BP3\|FAM108A9P\|RP11-652G5.2\|RP11-56L13.7\|RP11-1292F20.1\|RP11-652G5.1 |
| 16 | 16 | 32707001 | 32784000 | 76999 | DUP | Het | RP11-586K12.5\|HERC2P5\|RP11-586K12.4\|RP11-586K12.7\|FAM108A8P |
| 17 | 17 | 21321279 | 21322820 | 1541 | BND | Het | KCNJ12 |
| 17 | 17 | 72728980 | 72729594 | 614 | BND | Het | RAB37 |
| 18 | 18 | 12094507 | 12094672 | 165 | DEL | Het | RP11-815J4.5\|ANKRD62\|RNU6-324P |
| 18 | 18 | 50488067 | 50489494 | 1427 | BND | Het | DCC |
| 19 | 19 | 14944702 | 14944808 | 106 | BND | Het | OR7A5 |
| 20 | 20 | 135121 | 136537 | 1416 | BND | Het | DEFB127 |
| 20 | 20 | 26209794 | 26210981 | 1187 | DUP | Hom | MIR663A |

| | | | | | | | |
|---|---|---|---|---|---|---|---|
| 20 | 20 | 61347852 | 61347968 | 116 | BND | Het | NTSR1 |
| 22 | 22 | 21737001 | 21743000 | 5999 | DEL | Het | SCARNA18\|SCARNA17\|RIMBP3B\|RN7SKP63 |
| X | X | 2548047 | 2548362 | 315 | INV | Het | CD99P1 |
| X | X | 52263001 | 52567000 | 303999 | DEL | Het | XAGE1A\|XAGE1C\|XAGE2\|RP11-472D17.1\|RP11-472D17.6\|RP11-204I15.1\|RBM22P10\|RBM22P11\|RP11-472D17.4\|RP11-472D17.3\|RP11-472D17.2\|RBM22P8\|RBM22P9\|XAGE1D\|XAGE1E |
| X | X | 52886897 | 52887007 | 110 | BND | Het | XAGE3 |
| X | X | 58442001 | 58455000 | 12999 | DUP | Het | |
| X | X | 134960001 | 134965000 | 4999 | DEL | Het | CT45A6 |
| X | X | 143236001 | 143267000 | 30999 | DEL | Het | |
| X | X | 145891001 | 145896000 | 4999 | DEL | Het | CXorf51B\|CXorf51A |
| X | X | 148616587 | 148616693 | 106 | BND | Het | LINC00893\|IDS |
| X | X | 151082001 | 151088000 | 5999 | DEL | Het | RP11-366F6.2\|MAGEA4 |
| X | X | 154715001 | 154730000 | 14999 | DEL | Het | TMLHE-AS1\|RP11-218L14.4\|TMLHE |
| Y | Y | 58892001 | 58968000 | 75999 | DEL | Het | |
| MT | MT | 1 | 16568 | 16567 | BND | Hom | MT-ND1\|MT-ND2\|MT-TL1\|MT-TQ\|MT-TV\|MT-RNR1\|MT-RNR2\|MT-TM\|MT-TI\|MT-TF |
| GL000213.1 | GL000213.1 | 100001 | 105000 | 4999 | DEL | Het | MIR3118-5\|BX072566.1 |
| GL000205.1 | GL000205.1 | 144047 | 144162 | 115 | DEL | Het | AC011841.9\|AC011841.10 |
| GL000192.1 | GL000192.1 | 426001 | 437000 | 10999 | DEL | Het | |

Structural Variants

| ChromosomeA | chromosomeB | PosA | PosB | Length | variant | P10051_104 | Genes |
|---|---|---|---|---|---|---|---|
| 1 | 1 | 13138001 | 13143000 | 4999 | DEL | Het | RP13-221M14.2\|PRAMEF25 |
| 1 | 1 | 13350001 | 13358000 | 7999 | DEL | Het | PRAMEF5\|RP11-248D7.2 |
| 1 | 1 | 13369001 | 13374000 | 4999 | DEL | Het | PRAMEF5 |
| 1 | 1 | 108979001 | 108984000 | 4999 | DEL | Het | NBPF6\|RP11-131J3.1 |
| 1 | 1 | 121132677 | 206583277 | 85450600 | BND | Het | RP11-343N15.1\|RP11-343N15.5\|SRGAP2C |
| 1 | 1 | 145092950 | 145097082 | 4132 | BND | Het | SEC22B\|RP4-725K1.1 |
| 1 | 1 | 148781001 | 148786000 | 4999 | DEL | Het | RP11-763B22.10 |
| 1 | 1 | 148849596 | 148849731 | 135 | DUP | Hom | RP11-763B22.9\|RP11-763B22.6\|RP11-763B22.7 |
| 1 | 1 | 182420489 | 182420608 | 119 | BND | Het | RGSL1 |
| 1 | 1 | 226089867 | 226090915 | 1048 | DEL | Het | RP4-559A3.7\|LEFTY1 |
| 2 | 2 | 1533643 | 1535348 | 1705 | BND | Het | TPO |
| 2 | 2 | 34696001 | 34704000 | 7999 | DEL | Het | AC073218.1 |
| 2 | 2 | 86950001 | 86956000 | 5999 | DEL | Het | CHMP3\|RMND5A\|RNF103-CHMP3 |
| 2 | 2 | 92207001 | 92326000 | 118999 | DUP | WT | IGKV1OR2-2 |
| 2 | 2 | 132977817 | 132980438 | 2621 | BND | Het | ANKRD30BL |
| 2 | 2 | 181988057 | 181988239 | 182 | BND | Het | AC104820.2 |
| 4 | 4 | 49570001 | 49596000 | 25999 | DUP | WT | AC119751.1\|AC119751.2\|AC119751.3\|AC119751.5\|SNX18P25\|RP11-241F15.10 |
| 4 | 4 | 188594196 | 188594328 | 132 | BND | Het | RP11-565A3.2 |
| 5 | 5 | 68929001 | 69733000 | 803999 | DEL | Het | SMN2\|RP11-974F13.5\|RP11-98J23.1\|SERF1B\|RP11-1415C14.2\|RN7SL9P\|RP11-1319K7.1\|GTF2H2B\|RP11-1415C14.4\|CDH12P3\|RP11-848G14.5\|RP11-1415C14.1\|GUSBP3\|RP11-1415C14.3\|CDH12P2\|RP11-98J23.2 |
| 5 | 5 | 177156001 | 177162000 | 5999 | DEL | Het | FAM153A |
| 5 | 5 | 177221001 | 177227000 | 5999 | DEL | Het | RP11-1026M7.3\|RP11-1026M7.2 |
| 6 | 6 | 57284913 | 57289357 | 4444 | BND | Het | PRIM2 |
| 6 | 6 | 57423088 | 57429120 | 6032 | BND | Het | PRIM2 |
| 7 | 7 | 74309001 | 74314000 | 4999 | DEL | Het | Y_RNA\|STAG3L2\|PMS2P5 |
| 7 | 7 | 74561001 | 74572000 | 10999 | DUP | Het | GTF2IRD2B\|NCF1C |
| 7 | 7 | 133575001 | 133580000 | 4999 | DEL | Het | EXOC4 |
| 8 | 8 | 7418001 | 7425000 | 6999 | DEL | Het | FAM90A21P\|FAM90A22P\|FAM90A7P |
| 8 | 8 | 64711001 | 64716000 | 4999 | DEL | Het | RP11-32K4.1 |
| 8 | 8 | 142256585 | 142256707 | 122 | BND | Het | SLC45A4 |
| 9 | 9 | 78001 | 85000 | 6999 | DEL | Het | RP11-143M1.2\|RP11-143M1.7 |

| | | | | | | | |
|---|---|---|---|---|---|---|---|
| 9 | 9 | 40703001 | 40709000 | 5999 | DEL | Het | RP11-395E19.2\|SPATA31A3\|RP11-395E19.5 |
| 9 | 9 | 45544001 | 45574000 | 29999 | DEL | Het | RP11-187C18.3 |
| 10 | 10 | 14761833 | 14761951 | 118 | BND | Het | FAM107B\|RP11-398C13.2 |
| 10 | 10 | 47724001 | 47729000 | 4999 | DEL | Het | CTGLF11P |
| 10 | 10 | 47950001 | 47956000 | 5999 | DEL | Het | FAM21B |
| 10 | 10 | 48664001 | 48804000 | 139999 | DEL | Het | PTPN20B |
| 10 | 10 | 48807001 | 48841000 | 33999 | DEL | Het | FRMPD2P1\|PTPN20B |
| 10 | 10 | 49268001 | 49276000 | 7999 | DEL | Het | PTPN20CP\|BMS1P7 |
| 10 | 10 | 49277001 | 49333000 | 55999 | DEL | Het | PTPN20CP |
| 10 | 10 | 49373001 | 49378000 | 4999 | DEL | Het | FRMPD2 |
| 10 | 10 | 52510001 | 52516000 | 5999 | DEL | Het | ASAH2B |
| 10 | 10 | 115449672 | 115449800 | 128 | BND | Het | CASP7 |
| 11 | 11 | 48367530 | 48373694 | 6164 | BND | Het | OR4C4P |
| 11 | 11 | 95574875 | 95578117 | 3242 | BND | Het | RNA5SP345\|MTMR2 |
| 11 | 11 | 95595269 | 95595435 | 166 | BND | Het | MTMR2 |
| 11 | 11 | 114433315 | 131230465 | 16797150 | BND | Het | NXPE1 |
| 12 | 12 | 21185611 | 21194480 | 8869 | DEL | Het | SLCO1B3\|LST3\|SLCO1B7 |
| 12 | 12 | 50292024 | 50292132 | 108 | BND | Het | FAIM2 |
| 12 | 12 | 104278731 | 104278869 | 138 | BND | Het | RP11-642P15.1 |
| 12 | 12 | 110997307 | 111001268 | 3961 | DUP | Het | PPTC7 |
| 15 | 15 | 28611001 | 28617000 | 5999 | DEL | Het | RP11-483E23.3 |
| 15 | 15 | 28792001 | 28822000 | 29999 | DUP | Het | RP11-578F21.4\|RP11-483E23.7\|RP11-665A22.1\|RP11-536P16.4 |
| 15 | 15 | 30596001 | 30881000 | 284999 | DEL | Het | AC019322.1\|U8\|DNM1P50\|RP11-382B18.5\|RN7SL196P\|GOLGA8Q\|GOLGA8R\|AC026150.2\|Y_RNA\|RP11-382B18.3\|RP11-382B18.4\|CTD-3092A11.1\|CTD-3092A11.2\|AC026150.6\|AC026150.8\|CHRFAM7A\|ULK4P2\|RN7SL796P |
| 15 | 15 | 32729001 | 32759000 | 29999 | DEL | Het | AC135983.1\|RP11-632K20.8\|DNM1P32\|GOLGA8O\|ULK4P1\|RN7SL539P\|RP13-395E19.3 |
| 15 | 15 | 34702001 | 34708000 | 5999 | DEL | Het | GOLGA8A\|RP11-1H8.1 |
| 15 | 15 | 75867352 | 75867467 | 115 | DUP | Hom | PTPN9\|CTD-2323K18.1 |
| 15 | 15 | 85733001 | 85741000 | 7999 | DEL | Het | CSPG4P12 |
| 15 | 15 | 102511001 | 102522000 | 10999 | DUP | WT | DDX11L9\|WASH3P |
| 16 | 16 | 2669001 | 2675000 | 5999 | DEL | Het | PDPK2\|AC141586.5 |
| 16 | 16 | 78757422 | 78757547 | 125 | BND | Het | WWOX |

| | | | | | | | |
|---|---|---|---|---|---|---|---|
| 17 | 17 | 20410001 | 20417000 | 6999 | DEL | Het | KRT16P3\|AC025627.9\|RP11-434D2.7 |
| 17 | 17 | 21321279 | 21322822 | 1543 | BND | Het | KCNJ12 |
| 17 | 17 | 60807286 | 60807431 | 145 | BND | Het | RP11-156L14.1\|MARCH10 |
| 18 | 18 | 11471001 | 11611000 | 139999 | DUP | Het | RP11-128P17.3\|RP11-677O4.3\|RP11-712C7.1\|RP11-712C7.2 |
| 18 | 18 | 11471291 | 11880723 | 409432 | DUP | Het | RP11-78A19.4\|RP11-78A19.3\|RP11-78A19.2\|RP11-128P17.3\|RP11-677O4.2\|RP11-677O4.3\|RP11-677O4.4\|RP11-677O4.5\|RP11-677O4.6\|RP11-677O4.7\|RP11-712C7.1\|NPIPB1P\|RP11-712C7.2\|GNAL\|CHMP1B\|MPPE1 |
| 18 | 18 | 11646001 | 11881000 | 234999 | DUP | Het | RP11-78A19.4\|RP11-78A19.3\|RP11-78A19.2\|RP11-677O4.2\|RP11-677O4.4\|RP11-677O4.7\|GNAL\|CHMP1B\|MPPE1 |
| 18 | 18 | 20563631 | 20563763 | 132 | BND | Het | RBBP8 |
| 18 | 18 | 50488067 | 50489494 | 1427 | BND | Het | DCC |
| 18 | 18 | 54493289 | 54493508 | 219 | BND | Het | WDR7 |
| 19 | 19 | 2247066 | 2247173 | 107 | BND | Het | AMH\|MIR4321\|SF3A2 |
| 19 | 19 | 5761476 | 5761631 | 155 | BND | Het | CATSPERD |
| 19 | 19 | 20508001 | 20513000 | 4999 | DEL | Het | ZNF826P\|MIR1270-1 |
| 20 | 20 | 135121 | 136537 | 1416 | BND | Het | DEFB127 |
| 20 | 20 | 26209794 | 26210981 | 1187 | DUP | Hom | MIR663A |
| 20 | 20 | 41512475 | 41512592 | 117 | DUP | Het | PTPRT |
| 22 | 22 | 16598001 | 16615000 | 16999 | DUP | Het | |
| 22 | 22 | 20239138 | 20239432 | 294 | DUP | Hom | MIR1286\|RTN4R |
| 22 | 22 | 20366001 | 20500000 | 133999 | DEL | Het | GGTLC3\|snoU13\|RIMBP3\|AC023490.1\|AC023490.2\|SCARNA17\|PPP1R26P2\|RN7SKP131\|XXbac-B33L19.10\|CA15P2\|SCARNA18\|PI4KAP1 |
| 22 | 22 | 45677910 | 45678081 | 171 | BND | Het | UPK3A |
| X | X | 289001 | 294000 | 4999 | DEL | Het | PPP2R3B |
| X | X | 3748001 | 3832000 | 83999 | DEL | Het | RP11-706O15.5\|RP11-706O15.1\|RP11-706O15.3 |
| X | X | 3837001 | 3844000 | 6999 | DEL | Het | RP11-706O15.5\|RP11-706O15.7 |
| X | X | 48240001 | 48246000 | 5999 | DEL | Het | SSX4\|RP11-344N17.12 |
| X | X | 49190001 | 49209000 | 18999 | DUP | Het | GAGE2D\|GAGE2E\|GAGE13\|GAGE12J |
| X | X | 52517001 | 52527000 | 9999 | DEL | Het | XAGE1C\|XAGE1D\|RBM22P9 |
| X | X | 52553001 | 52565000 | 11999 | DEL | Het | RBM22P11 |
| X | X | 143236001 | 143247000 | 10999 | DEL | Het | |
| X | X | 151080001 | 151088000 | 7999 | DEL | Het | RP11-366F6.2\|MAGEA4 |
| X | X | 154715001 | 154721000 | 5999 | DEL | Het | TMLHE-AS1\|RP11-218L14.4\|TMLHE |
| X | X | 155261001 | 155271000 | 9999 | DUP | Het | DDX11L16 |

| | | | | | | | |
|---|---|---|---|---|---|---|---|
| MT | MT | 1 | 16568 | 16567 | BND | Hom | MT-ND1\|MT-ND2\|MT-TL1\|MT-TQ\|MT-TV\|MT-RNR1\|MT-RNR2\|MT-TM\|MT-TI\|MT-TF |
| GL000227.1 | GL000227.1 | 102001 | 129000 | 26999 | DEL | Het | |
| GL000220.1 | GL000220.1 | 104132 | 104251 | 119 | DUP | Hom | AL592188.2\|AL592188.3\|AL592188.8\|AL592188.4\|RNA18S5 |
| GL000200.1 | GL000200.1 | 19001 | 34000 | 14999 | DEL | Het | |
| GL000200.1 | GL000200.1 | 35001 | 188000 | 152999 | DEL | Het | |

Structural Variants

| ChromosomeA | chromosomeB | PosA | PosB | Length | variant | P10051_105 | Genes |
|---|---|---|---|---|---|---|---|
| 1 | 1 | 12856001 | 12865000 | 8999 | DEL | Het | PRAMEF1 |
| 1 | 1 | 13028001 | 13033000 | 4999 | DEL | Het | PRAMEF22\|PRAMEF6 |
| 1 | 1 | 13138001 | 13143000 | 4999 | DEL | Het | RP13-221M14.2\|PRAMEF25 |
| 1 | 1 | 13403001 | 13410000 | 6999 | DEL | Het | RP11-219C24.6 |
| 1 | 1 | 143136001 | 143198000 | 61999 | DUP | WT | RP11-782C8.4\|RP11-782C8.2\|RP11-782C8.1\|RP11-782C8.3\|MIR3118-2 |
| 1 | 1 | 143217001 | 143244000 | 26999 | DUP | Het | RP11-782C8.5\|RP11-782C8.7\|RP11-782C8.6\|RP11-782C8.1\|RP11-782C8.8 |
| 1 | 1 | 143370001 | 143500000 | 129999 | DUP | WT | RP11-435B5.4\|RP11-435B5.3\|RP11-435B5.6\|BX004987.1\|MIR3118-3\|RP11-435B5.5 |
| 1 | 1 | 144283001 | 144496000 | 212999 | DEL | Het | LINC00623\|AL592284.1\|RP11-640M9.1\|RP6-137J22.3\|PPIAL4B\|RNVU1-4\|RNVU1-5 |
| 1 | 1 | 145092950 | 145097082 | 4132 | BND | Het | SEC22B\|RP4-725K1.1 |
| 1 | 1 | 148849594 | 148849728 | 134 | DUP | Hom | RP11-763B22.9\|RP11-763B22.6\|RP11-763B22.7 |
| 1 | 1 | 171848819 | 171863952 | 15133 | DEL | Het | DNM3 |
| 1 | 1 | 246243534 | 246243651 | 117 | BND | Het | SMYD3 |
| 2 | 2 | 71890068 | 71890172 | 104 | BND | Het | DYSF |
| 2 | 2 | 89569001 | 89575000 | 5999 | DEL | Het | IGKV3-34\|IGKV1-33 |
| 2 | 9 | 156528084 | 128068328 | #NUM! | BND | Het | |
| 2 | 2 | 236907288 | 236907419 | 131 | BND | Het | AGAP1 |
| 3 | 3 | 151675618 | 151693804 | 18186 | DEL | Het | |
| 3 | 3 | 197826001 | 197833000 | 6999 | DEL | Het | AC073135.3 |
| 21 | 4 | 34205582 | 59654 | #NUM! | BND | Het | C21orf49\|ZNF595 |
| 4 | 4 | 7512270 | 7512457 | 187 | DUP | Het | SORCS2 |
| 4 | 4 | 49570001 | 49594000 | 23999 | DUP | WT | AC119751.1\|RP11-241F15.10\|SNX18P25\|AC119751.5 |
| 12 | 4 | 109544728 | 112495903 | #NUM! | BND | Het | \|RP11-968O1.5\|UNG |
| 4 | 4 | 146613018 | 146613127 | 109 | BND | Het | C4orf51 |
| 11 | 4 | 51376183 | 168319545 | #NUM! | BND | Het | \|RN7SL776P |
| 4 | 4 | 170708287 | 170708397 | 110 | DUP | Het | AC106878.1\|PTGES3P3 |
| 4 | 4 | 178657744 | 178657848 | 104 | BND | Het | LINC01099\|LINC01098 |
| 5 | 8 | 167285630 | 126483500 | #NUM! | BND | Het | TENM2 |
| 5 | 5 | 177154001 | 177162000 | 7999 | DEL | Het | FAM153A |
| 5 | 5 | 179934820 | 179935072 | 252 | BND | Het | CNOT6 |
| 6 | 6 | 31998001 | 32004000 | 5999 | DEL | Het | CYP21A2\|TNXB\|C4B\|C4B-AS1 |
| 6 | 6 | 57284913 | 57289357 | 4444 | BND | Het | PRIM2 |
| 6 | 6 | 57423088 | 57429120 | 6032 | BND | Het | PRIM2 |
| 7 | 7 | 72640001 | 72680000 | 39999 | DEL | Het | NCF1B\|GTF2IRD2P1 |
| 7 | 7 | 143907001 | 143912000 | 4999 | DEL | Het | ARHGEF35\|RP4-545C24.1 |
| 7 | 7 | 143989001 | 143994000 | 4999 | DEL | Het | OR2A9P\|OR2A1-AS1\|ARHGEF35\|ARHGEF34P |

| | | | | | | | |
|---|---|---|---|---|---|---|---|
| 8 | 8 | 92001 | 99000 | 6999 | DEL | Het | RP11-585F1.8 |
| 8 | 8 | 7617001 | 7622000 | 4999 | DEL | Het | FAM90A17P\|FAM90A19P\|FAM90A10P\|FAM90A9P |
| 8 | 8 | 43586201 | 43604319 | 18118 | BND | Het | |
| 8 | 8 | 110077365 | 110077475 | 110 | BND | Het | RP11-1084E5.1 |
| 8 | 8 | 145290001 | 145446000 | 155999 | DEL | Hom | SCXB\|KM-PA-2\|MROH1\|FAM203B |
| 8 | 8 | 145448001 | 145501000 | 52999 | DEL | Het | AC110280.1\|CTD-3232M19.2\|BOP1\|SCXA |
| 9 | 9 | 39185001 | 40147000 | 961999 | DEL | Het | RN7SL462P\|CTD-2173L22.4\|CNTNAP3\|SPATA31A1\|VN2R4P\|SPATA31A2\|RP11-381G8.1\|FAM74A5\|BX664726.3\|FAM74A1\|RP11-347J14.7\|RP11-347J14.4\|AL353791.1\|RP11-347J14.8\|ATP5A1P9\|BX088645.3\|BX088645.2\|RN7SL640P\|RP11-133G22.1\|RP11-95H8.5\|AL590812.1\|AL590812.3 |
| 9 | 9 | 40148001 | 40160000 | 11999 | DEL | Het | |
| 9 | 9 | 40381001 | 40399000 | 17999 | DEL | Het | VN2R5P |
| 9 | 9 | 41824001 | 41844000 | 19999 | DEL | Het | |
| 9 | 9 | 43089001 | 43096000 | 6999 | DEL | Het | ANKRD20A3 |
| 9 | 9 | 66344001 | 66350000 | 5999 | DEL | Het | CNN2P5\|RP11-459O16.1 |
| 9 | 9 | 95190968 | 95191118 | 150 | BND | Het | CENPP\|OMD |
| 10 | 10 | 6837001 | 6843000 | 5999 | DEL | Het | LINC00707 |
| 10 | 10 | 11774001 | 11780000 | 5999 | DEL | Het | ECHDC3 |
| 10 | 10 | 47715001 | 48002000 | 286999 | DEL | Het | SLC9A3P4\|FAM25HP\|ASAH2C\|ANXA8L2\|AL603965.1\|CTGLF11P\|FAM21B |
| 10 | 10 | 52509001 | 52516000 | 6999 | DEL | Het | ASAH2B |
| 10 | 10 | 54287637 | 78377068 | 2E+07 | BND | Het | |
| 10 | 10 | 66017499 | 66037818 | 20319 | DEL | Het | |
| 11 | 13 | 38232929 | 64492165 | #NUM! | BND | Het | RP11-436H16.1 |
| 11 | 11 | 48367530 | 48373694 | 6164 | BND | Het | OR4C4P |
| 11 | 11 | 48889052 | 48900530 | 11478 | DUP | Hom | RP11-56P9.5\|RP11-56P9.10 |
| 11 | 11 | 68859854 | 68859986 | 132 | DUP | Het | TPCN2 |
| 11 | 11 | 75676436 | 75685566 | 9130 | DEL | Het | UVRAG |
| 11 | 11 | 85149976 | 85177053 | 27077 | DEL | Het | DLG2\|RP11-482L11.1 |
| 11 | 11 | 114433315 | 131230467 | 2E+07 | BND | Het | NXPE1 |
| 12 | 12 | 31272660 | 31272790 | 130 | BND | Het | RP11-551L14.1 |
| 12 | 12 | 50292024 | 50292132 | 108 | BND | Het | FAIM2 |
| 12 | 12 | 54147589 | 54147701 | 112 | BND | Het | RP11-686F15.2\|RP11-686F15.3 |
| 14 | 18 | 26878436 | 26463996 | #NUM! | BND | Het | |
| 14 | 14 | 39502719 | 39502847 | 128 | BND | Het | SEC23A |
| 14 | 14 | 69780314 | 69780484 | 170 | BND | Het | GALNT16 |
| 14 | 14 | 72310001 | 72316000 | 5999 | DEL | Het | RP6-114E22.1 |

| | | | | | | | |
|---|---|---|---|---|---|---|---|
| 15 | 15 | 21050001 | 21110000 | 59999 | DUP | WT | RP11-810K23.9\|RP11-810K23.10\|RP11-810K23.5\|POTEB2\|RNU6-749P |
| 15 | 15 | 28611001 | 28617000 | 5999 | DEL | Het | RP11-483E23.3 |
| 15 | 15 | 28829001 | 28866000 | 36999 | DUP | Het | HERC2P9\|RP11-665A22.1 |
| 15 | 20 | 89566414 | 33736945 | #NUM! | BND | Het | |
| 16 | 16 | 29467001 | 29503000 | 35999 | DUP | Het | SNX29P2\|RP11-231C14.3\|SULT1A4\|SLX1B\|RP11-345J4.1\|RP11-345J4.3\|RP11-345J4.5\|SLX1B-SULT1A4\|BOLA2\|RP11-231C14.4\|RP11-231C14.5 |
| 16 | 16 | 32944001 | 33642000 | 697999 | DUP | WT | IGHV1OR16-2\|RP11-1277H1.3\|RP11-293B20.3\|FAM108A10P\|TP53TG3B\|TP53TG3C\|IGHV3OR16-8\|RP11-989E6.8\|RP11-19N8.2\|RP11-19N8.3\|RP11-19N8.4\|RP11-19N8.6\|RP11-19N8.7\|HERC2P8\|RP11-104C4.2\|RP11-104C4.4\|BMS1P8\|RP11-989E6.10\|RP11-989E6.11\|RP11-1437A8.4\|RP11-1437A8.5\|RP11-1437A8.6\|RP11-1437A8.7\|RP11-293B20.2\|RP11-1437A8.2\|RP11-1437A8.3\|RP11-23E10.2\|RP11-23E10.3\|RP11-23E10.4\|RP11-23E10.5\|IGHV3OR16-13\|IGHV3OR16-12\|IGHV3OR16-10\|ENPP7P13\|IGHV1OR16-4 |
| 16 | 16 | 33685001 | 33789000 | 103999 | DUP | Het | IGHV3OR16-7\|RP11-812E19.5\|RP11-812E19.14\|RP11-812E19.3\|RP11-598D12.4\|RP11-598D12.3\|ARHGAP23P1\|IGHV3OR16-16 |
| 17 | 17 | 1141582 | 1141730 | 148 | BND | Het | AC144836.1 |
| 17 | 17 | 10563731 | 10580449 | 16718 | DUP | Het | CTC-297N7.1\|MYH3\|CTC-297N7.10\|SCO1\|CTC-297N7.8 |
| 17 | 17 | 21321279 | 21322822 | 1543 | BND | Het | KCNJ12 |
| 17 | 17 | 44598001 | 44605000 | 6999 | DEL | Het | LRRC37A2\|ARL17A |
| 17 | 17 | 76393372 | 76393475 | 103 | BND | Het | SNORA30\|PGS1 |
| 17 | 17 | 79399740 | 79399873 | 133 | BND | Hom | RP11-1055B8.7 |
| 18 | 18 | 50488067 | 50489494 | 1427 | BND | Het | DCC |
| 19 | 19 | 6995404 | 6995509 | 105 | BND | Het | EMR4P |
| 19 | 19 | 9246113 | 9247205 | 1092 | DEL | Het | ZNF317 |
| 19 | 19 | 50517687 | 50517795 | 108 | BND | Het | VRK3 |
| 20 | 20 | 135121 | 136537 | 1416 | BND | Het | DEFB127 |
| 21 | 21 | 10382001 | 10474000 | 91999 | DUP | WT | SNORA70\|RN7SL52P\|bP-21201H5.1 |
| 21 | 21 | 10476001 | 10648000 | 171999 | DUP | WT | bP-21201H5.1 |
| X | X | 330001 | 748000 | 417999 | DUP | WT | PPP2R3B\|SHOX\|AL732314.1\|KRT18P53\|FABP5P13 |
| X | X | 1438187 | 1438648 | 461 | BND | Het | RN7SL355P |
| X | X | 24671988 | 24672136 | 148 | BND | Het | PCYT1B-AS1\|PCYT1B |
| X | X | 52886896 | 52887008 | 112 | BND | Het | XAGE3 |
| X | X | 103150001 | 103155000 | 4999 | DEL | Hom | RP11-370B6.1 |
| X | X | 134942001 | 134948000 | 5999 | DEL | Het | CT45A5\|CT45A4 |
| X | X | 151081001 | 151086000 | 4999 | DEL | Het | RP11-366F6.2\|MAGEA4 |
| X | X | 154580001 | 154585000 | 4999 | DEL | Het | RP13-228J13.5\|RP13-228J13.1 |
| Y | Y | 10011001 | 10025000 | 13999 | DUP | WT | CDC27P2\|AC006987.6\|AC006987.7\|PCMTD1P1 |

| | | | | | | | |
|---|---|---|---|---|---|---|---|
| MT | MT | 1 | 16568 | 16567 | BND | Hom | MT-ND1\|MT-ND2\|MT-TL1\|MT-TQ\|MT-TV\|MT-RNR1\|MT-RNR2\|MT-TM\|MT-TI\|MT-TF |
| GL000217.1 | GL000217.1 | 118001 | 161000 | 42999 | DUP | WT | |
| GL000194.1 | GL000194.1 | 82001 | 114000 | 31999 | DUP | WT | AC145212.3\|AC145212.2 |
